# Comparative genomic analyses and a novel linkage map for cisco (*Coregonus artedi*) provide insights into chromosomal evolution and rediploidization across salmonids

**DOI:** 10.1101/834937

**Authors:** Danielle M. Blumstein, Matthew A. Campbell, Matthew C. Hale, Ben J. G. Sutherland, Garrett J. McKinney, Wendylee Stott, Wesley A. Larson

## Abstract

Whole-genome duplication (WGD) is hypothesized to be an important evolutionary mechanism that can facilitate adaptation and speciation. Genomes that exist in states of both diploidy and residual tetraploidy are of particular interest, as mechanisms that maintain the ploidy mosaic after WGD may provide important insights into evolutionary processes. The Salmonidae family exhibits residual tetraploidy, and this, combined with the evolutionary diversity formed after an ancestral autotetraploidization event, makes this group a useful study system. In this study, we generate a novel linkage map for cisco (*Coregonus artedi*), an economically and culturally important fish in North America and a member of the subfamily Coregoninae, which previously lacked a high-density haploid linkage map. We also conduct comparative genomic analyses to refine our understanding of chromosomal fusion/fission history across salmonids. To facilitate this comparative approach, we use the naming strategy of protokaryotype identifiers (PKs) to associate duplicated chromosomes to their putative ancestral state. The female linkage map for cisco contains 20,292 loci, 3,225 of which are likely within residually tetraploid regions. Comparative genomic analyses revealed that patterns of residual tetrasomy are generally conserved across species, although interspecific variation persists. To determine the broad-scale retention of residual tetrasomy across the salmonids, we analyze sequence similarity of currently available genomes and find evidence of residual tetrasomy in seven of the eight chromosomes that have been previously hypothesized to show this pattern. This interspecific variation in extent of rediploidization may have important implications for understanding salmonid evolutionary histories and informing future conservation efforts.

## Introduction

The evolutionary significance of whole-genome duplications (WGDs) has been intensively debated for decades (e.g; Ohno 1970; Taylor *et al.* 2003; Santini *et al.* 2009; Wood *et al.* 2009; Zhan *et al.* 2014; Mayrose *et al.* 2015; Van de Peer *et al.* 2017). Multiple studies have hypothesized that WGD is an important evolutionary mechanism that can facilitate adaptation on short- and long-term evolutionary timescales (Ohta 1989, Selmecki *et al.* 2015; Van de Peer *et al.* 2017). For example, genes found in polyploid regions are able to gain new function (i.e., neofunctionalization) without the consequences of deleterious mutations affecting the main function of the original gene copy. This may facilitate adaptive molecular divergence and evolution of new phenotypes (Wittbrodt *et al.* 1998; Wendel 2000; Rastogi and Liberles 2005). However, other studies have hypothesized that WGD presents significant challenges for meiosis and mitosis (Hollister 2015) and may not have as much of an effect on evolution as originally considered (Mayrose *et al.* 2011; Arrigo and Barker 2012; Vanneste *et al.* 2014; Clarke *et al.* 2016). A consensus is therefore yet to be reached on the evolutionary impact of WGD relative to other evolutionary forces.

Conducting genetic studies on organisms with relatively recent WGD can be challenging due to the inability to differentiate alleles and sequences from the same chromosome (homologs) from those on the duplicated chromosome (homeologs) (Limborg *et al.* 2016b). Fortunately, approaches leveraging gamete manipulation, high sequencing coverage, and long read sequencing have improved our ability to characterize duplicated regions. Linkage mapping with haploids and doubled haploids has facilitated analysis of duplicated regions in salmonids (Brieuc *et al.* 2014; Kodama *et al.* 2014; Lien *et al.* 2016; Waples *et al.* 2016). Further, long-read sequencing technologies have made it possible to assemble complex genomes with convoluted duplication histories (Kyriakidou *et al.* 2018). These technological advances have revolutionized our ability to understand genomic architecture in species that have adaptively radiated into dozens of species following an ancestral WGD in lineages, such as salmonids (Lien *et al.* 2016; Robertson *et al.* 2017; Campbell *et al.* 2019) and many plant species (Alix *et al.* 2017).

Salmonids are derived from an ancestral species that underwent a WGD ~100 million years ago (Ss4R, Allendorf and Thorgaard 1984; Berthelot *et al.* 2014; Macqueen and Johnston 2014; Lien *et al.* 2016) and have since diversified into a broad array of ecologically and genetically distinct taxa. The Salmonidae family is comprised of three subfamilies: Salmoninae (salmon, trout, and char), Thymallinae (graylings), and Coregoninae (whitefish and ciscoes) (Norden 1961), and diversification of these subfamilies post-dates the Ss4R, occurring 40-50 million years ago (Campbell *et al.* 2013; Macqueen and Johnston 2014). Phylogenetic analysis has revealed that the majority of the salmonid genome returned to a diploid inheritance state prior to the divergence of the subfamilies (Robertson *et al.* 2017). However, the rediploidization process is still incomplete and approximately 20-25% of each salmonid genome still shows signals of tetrasomic inheritance (i.e., residual tetrasomy, or the recombination between homeologs that results in the exchange of alleles between homeologous chromosomes (Allendorf *et al.* 2015; Lien *et al.* 2016; Robertson *et al.* 2017)).

Early evidence for residual tetrasomy in the salmonids was identified using allozyme studies in experimental crosses (Allendorf and Danzmann 1997) and, more recently, by linkage maps, sequenced genomes, and sequence capture (Kodama *et al.* 2014; McKinney *et al.* 2017; Robertson *et al.* 2017; Christensen *et al.* 2018b; Pearse *et al.* 2019). High-density linkage maps that include both duplicated and non-duplicated markers have revealed that eight pairs of homeologous chromosomes repeatedly display evidence of residual tetraploidy despite independent fusion and fission events (Brieuc *et al.* 2014; Kodama *et al.* 2014; Sutherland *et al.* 2016). These chromosomes, often referred to as the “magic eight,” have been observed in linkage mapping studies of coho salmon *Oncorhynchus kisutch* (Kodama *et al.* 2014), Chinook salmon *O. tshawytscha* (Brieuc *et al.* 2014; McKinney *et al.* 2016; McKinney *et al.* 2019), pink salmon *O. gorbuscha* (Tarpey *et al.* 2017), chum salmon *O. keta* (Waples *et al.* 2016), and sockeye salmon *O. nerka* (Larson *et al.* 2015). Mapping studies have also revealed that at least one of the two homeologs exhibiting residual tetraploidy is within a chromosomal fusion, suggesting their role in tetrasomy persistence (Brieuc *et al.* 2014; Kodama *et al.* 2014; Sutherland *et al.* 2016). Analysis of sequenced genomes for Atlantic salmon (*Salmo salar*) and rainbow trout (*O. mykiss*) also support evidence of residual tetraploidy, although in these genomic studies only seven of these pairs were identified as displaying clear signals of conserved residual tetraploidy (Lien *et al.* 2016; Campbell *et al.* 2019). Sequence capture analysis also identified seven pairs displaying conserved signals of residual tetrasomy across species (Robertson *et al.* 2017). This led to the definition of two types of homeologous regions: 1) ancestral ohnologue resolution (AORe) regions with relatively low sequence similarity with ancestral homeologs that likely rediploidized prior to species diversification; and 2) lineage-specific ohnologue resolution (LORe) regions with high sequence similarity among homeologs, likely maintained by residual tetraploidy (Robertson *et al.* 2017).

Over the past decade, an extensive proliferation of genomic resources has occurred within salmonids. Currently, linkage maps that include duplicated regions are available for five *Oncorhynchus* species, and genome assemblies are available for grayling *Thymallus thymallus* (Savilammi *et al.* 2019), Atlantic salmon (Lien *et al.* 2016), Arctic char *Salvelinus alpinus* (Christensen *et al.* 2018b), rainbow trout (Pearse *et al.* 2019), and Chinook salmon (Christensen *et al.* 2018a). These resources permit investigation into the processes of rediploidization and residual tetrasomy across the salmonid family. There is however an underrepresentation of other lineages within the salmonid family, such as the Coregoninae subfamily (but see Gagnaire *et al.* 2013, and recently De-Kayne *et al.* 2020).

Our focal species for this manuscript was the North American cisco (*Coregonus artedi*). Cisco are a commercially, economically, and ecologically important species across northern North America. Additionally, these species are preyed upon by many apex predators and have historically represented an important trophic linkage in freshwater ecosystems, such as in the Laurentian Great Lakes (Eshenroder *et al.* 2016). Cisco also display extremely high phenotypic diversity, which has led to the definition of multiple forms based primarily on morphological evidence (Eshenroder *et al.* 2016; Koelz 1929; Yule *et al.* 2013). Recent environmental shifts and a renewed focus on conservation of native species has resulted in increased interest in restoring cisco in Laurentian Great Lakes and other inland lakes in the United States and Canada (Zimmerman and Krueger 2009; Eshenroder *et al.* 2016). Key to this restoration effort is understanding the relative roles of phenotypic plasticity and adaptive genetic diversity in shaping phenotypic diversity within cisco, and genomic tools and resourced are needed to address these important questions.

In the current study, we develop a high-density linkage map for cisco, the first haploid linkage map for the Coregoninae subfamily, and analyze existing genomic resources for other salmonids with the goal of investigating patterns of residual tetrasomy and chromosomal fusion and fission history across the salmonid family, with particular focus on the coregonines. Our results suggest that (1) interspecific variation in residual tetrasomy is greater than previously observed; (2) binary definitions of chromosome ploidy status may not adequately capture variation within and among species; (3) linkage maps and sequenced genomes identify slightly different patterns regarding residual tetrasomy; and (4) a large number of fissions and fusions are specific to the base of the Coregoninae subfamily and species-specific fusions within Coregoninae are rare. This study uses new and existing resources to conduct the most comprehensive analysis of residual tetrasomy across the salmonid phylogeny to date.

## Methods

### Experimental crosses for linkage mapping

Genotypes from four diploid families (n = 73, 81, 84, and 95) and three haploid families (n = 80, 111, 139) were used to build sex-specific linkage maps (Table S2). Diploid crosses were constructed from cisco collected in northern Lake Huron (45° 58’51.6” N-84 °19’40.8” W, USA) during spawning season (November 2015) by U. S. Fish and Wildlife Service crews using standardized gill net assessment methods. Gametes for haploid crosses were collected following the same methods, from the same location and month but in 2017. Gametes were extracted from mature fish and eggs were combined directly with sperm to produce diploid crosses or with sperm that had been irradiated with 300,000 μJ/cm2 UV light for two minutes to break down the DNA and produce haploid crosses. UV irradiation leaves the sperm intact so that the egg can be activated but no paternal genetic material is contributed (i.e., gynogenesis, Chourrout 1982), resulting in haploid embryos with maternal genetic material only. Crosses were made in the field and transported to the U. S. Geological Survey-Great Lakes Science Center, Ann Arbor, Michigan (USA) for rearing. Tissue samples (fin clips) were taken from adult parents, from offspring of the diploid crosses at age two, and from haploids approximately 50 days post fertilization. All samples were preserved in a combination of 95% ethanol and 5% EDTA and sent to the University of Wisconsin, Stevens Point Molecular Conservation Genetics Lab for processing. Laboratory and field collections were conducted under the auspices of the U.S. Fish and Wildlife Service and U.S. Geological Survey-Great Lakes Science Center and all necessary animal care and use protocols were filed by these agencies.

### DNA extraction and RAD-sequencing library preparation

DNA was extracted using DNeasy 96 Blood and Tissue Kits (Qiagen, Valencia, California) per the manufacturer’s instructions. Quality and quantity of the extracted genomic DNA was measured using the Quant-iT PicoGreen double-stranded DNA Assay (Life Technologies) with a plate reader (BioTek). To confirm ploidy of haploid samples, parents and offspring were genotyped using six polymorphic microsatellite loci known to occur at diploid sites developed by Angers *et al.* (1995), Patton *et al.* (1997), and Rogers *et al.* (2004), and individuals were classified as haploids if only a single allele was present at all loci. The probability of not detecting a diploid if a diploid was present is ~1.09% based on microsatellite heterozygosity in the parental population (unpublished data, Wendylee Stott).

Genomic DNA from diploids and confirmed haploids was prepared for RAD sequencing using the *Sbf*I restriction enzyme following the methods outlined in Ali *et al.* (2016) except shearing restriction digested DNA was done with NEBNext® dsDNA Fragmentase® (New England Biolabs, Inc) instead of sonication. DNA was then purified and indexed using NEBNext® Ultra™ DNA Library Prep Kit for Illumina® per the manufacturer’s instructions (New England Biolabs, Inc). Libraries were sequenced on a HiSeq4000 with paired end 150bp chemistry at the Michigan State Genomics Core Facility (East Lansing, MI).

### SNP discovery and genotyping

Quality filtering, SNP identification, and genotyping was conducted using *Stacks* v.2.2 (Rochette and Catchen 2017). First, samples were demultiplexed with *process_radtags* with flags -c, -q, -r, -t 140, --*bestrad*. Markers were discovered *de novo* and genotyped within individuals with *ustacks* (flags = -m 3, -M 5, -H --*max_locus_stacks* 4, --*model_type* bounded, --*bound_high* 0.05, --*disable-gapped*). A catalog of loci was created using a subset of the individuals (diploid parents = 8, haploid parents = 5, wild fish = 38, total cisco = 51) with *cstacks* (-n of 3, --*disable-gapped*). The 38 wild fish used in the catalog were collected from the same geographic area using the same collection methods as listed above and were included to search for a sex identification marker, which was unsuccessful (*data not shown*).

Putative loci within each individual fish were matched against the catalog with *sstacks* (flag = --*disable-gapped*), *tsv2bam* was used with only the forward reads to orient the data by SNP, and *gstacks* was used to combine genotypes across individuals. Only the forward reads from the paired-end data were used in *gstacks* due to variable read depth in reverse reads and thus less reliable genotyping. *gstacks* was also run separately with the forward and reverse reads using *tsv2bam* to assemble longer contigs for sequence alignment and annotation. Final genotype calls were output as VCF files with *populations* (flags = -r 0.75), with each family grouped as a separate population in the *popmap* sample interpretation file. VCFtools (Danecek *et al.* 2011) was used to identify and remove individuals from the study that were missing more than 30% of data.

Maximum likelihood-based methods developed by Waples *et al.* (2016) were used to identify loci that could be mapped in haploid crosses and to identify potentially duplicated loci. Custom Python scripts available on GitHub (Python Software Foundation version 2.7) (see *Data Availability*), were used to filter the haplotype VCF file output from the *populations* module to identify loci that could be mapped in the diploid families. Loci missing more than 25% of data and loci that were genotyped as heterozygous in both parents of diploid families (and therefore could not be reliably mapped) were removed (as in Larson *et al.* 2015). Individual genotypes were exported with the custom Python scripts as *LepMap3* input files. As a final step before linkage mapping, genotypes from all seven families (haploid and diploid) were combined into a single dataset to form the final female *LepMap3* input file and the four diploid families were combined into a single dataset to form the final male *LepMap3* input file.

### Linkage mapping

The program *LepMap3* (Rastas 2017) was used to construct linkage maps following the methods of McKinney *et al.* (2016). Due to heterochiasmy (i.e., recombination rate differences between males and females) that occurs in the Salmonidae family (Sakamoto *et al.* 2000), a separate map was constructed for each sex. Loci were filtered and clustered into linkage groups (LGs) based on recombination rates by calculating logarithm of the odds (LOD) scores between all pairs of loci with the *SeparateChromosomes2* module. The LOD scores were chosen by increasing the LOD value by one with no minimum marker parameter until the number of LGs stabilized and was similar to that expected based on the haploid karyotype of cisco (N=40, Phillips *et al.* 1996) and the number of makers for additional LGs in the female map was less then 100 markers and less then 10 makers on the male map. The final LOD scores used to generate the map were LOD = 15 and 5 for the female and the male maps, respectively. Loci were then ordered within LGs by utilizing paternal and maternal haplotypes as inheritance vectors with the *OrderMarkers2* module. We used a minimum marker number per LG of 100 for the female map and 40 for the male map as LGs with fewer markers did not display consistent synteny with genomic resources (i.e. markers aligned to ≫2 chromosome arms) and were likely statistical artefacts (data not shown). LGs were reordered and markers removed until no large gaps remained (Rastas 2017).

### Comparative analysis of syntenic regions of linkage maps via MAPCOMP

MAPCOMP can be used to compare syntenic relationships among markers between linkage maps of any related species using a genome intermediate from another related species (Sutherland *et al.* 2016). Here, MAPCOMP was used to compare the cisco map with other *Coregonus* spp., including lake whitefish *C. clupeaformis* (Gagnaire *et al.* 2013), European whitefish *C. lavaretus* “Albock” (De-Kayne and Feulner 2018), as well as other representative species from other salmonid genera (i.e., Atlantic salmon (Lien *et al.* 2016), brook trout *S. fontinalis* (Sutherland *et al.* 2016), and Chinook salmon (Brieuc *et al.* 2014) and a representative outgroup to the salmonid WGD, northern pike *Esox lucius* (Rondeau *et al.* 2014). All code to collect and prepare maps, and run the analysis are available on GitHub (see *Data Availability*).

MapComp pairs loci between the two compared linkage maps if they align at the same locus or close to each other on the same contig or scaffold on the intermediate reference genome (Sutherland *et al.* 2016). Due to the large phylogenetic distance covered in this analysis, two reference genomes were used, including grayling (Savilammi *et al.* 2019) for comparisons within Coregonus and Atlantic salmon (Lien *et al.* 2016) for comparisons between all species. As salmonid chromosomal evolution is typified by Robertsonian fusions (Phillips and Rab 2001), fused chromosome arms in cisco, lake whitefish, and European whitefish were identified by aligning cisco markers to multiple salmonid genomes to identify cases where one cisco LG corresponded with at least two chromosome arms in another species. The fusion and fission phylogenetic history was plotted based on the most parsimonious explanation of common fusions among Coregonus spp., basing the approximate occurrences of fusions on shared fusions among species. Fusion history shared at the base of the Salmoninae lineage was taken from earlier work (Sutherland *et al.* 2016).

### Homeolog identification, similarity and inheritance mode

Homeologous chromosome arms can be identified in haploid crosses by mapping multiple alleles of duplicated markers based on the expected segregation ratio per paralog as described in Brieuc *et al.* (2014). Duplicated markers in cisco were mapped using this method, and duplicated markers from previously constructed linkage maps for coho salmon (Kodama *et al.* 2014), Chinook salmon (McKinney *et al.* 2019), pink salmon (Tarpey *et al.* 2017), chum salmon (Waples *et al.* 2016), and sockeye salmon (Larson *et al.* 2015) were obtained. Homeologs were then ranked based on the number of markers supporting each known homeologous relationship and the number of duplicated markers was used to determine patterns of inheritance (i.e., disomy vs. tetrasomy) of the homeologous pair.

Homeology was assessed by comparing DNA sequence similarity between homeologous arms from chromosome-level genome assemblies. Genomes included in this analysis were grayling (GCA_004348285.1, Savilammi *et al.* 2019), Atlantic salmon (GCF_000233375.1, Lien *et al.* 2016), Arctic char (GCF_002910315.2, Christensen *et al.* 2018b), rainbow trout (GCA_002163495.1, Pearse *et al.* 2019), and Chinook salmon (GCF_002872995.1, Christensen *et al.* 2018a). Homeologous arms were inferred either as identified in the original genome paper (references above) or through MapComp comparisons. Homeologous arms were then aligned to determine sequence similarity using LASTZ v1.02 (Harris 2007) following methods outlined in Lien *et al.* (2016). Options specified with LASTZ included --chain --gapped --gfextend -- identity=75.0‥100. --ambiguous=iupac --exact=20. The analysis was restricted to alignments with minimum percent match values of 75%, and a minimum length of 1,000 base pairs to minimize the likelihood of spurious alignments that might be due to gene family duplication rather than WGD. Overall similarity of a homeologous pair was represented by the median percent similarity of all alignments, weighted by alignment length, and summarized with boxplots for each homeologous pair in each species ordered based on descending median percentage sequence similarity.

Each homeologous pair was classified into one of two categories, tetrasomic or disomic, using a machine learning approach. Previous research indicates that salmonids are undergoing rediploidization of tetrasomic homeologous pairs, disomic homeologous pairs, and intermediate homeologous pairs of uncertain affinity (Campbell *et al.* 2019; Lien *et al.* 2016). To objectively classify protokaryotypes into tetrasomic homeologous or disomic homeologous pairs in each species, a training set was constructed containing the four highest and four lowest sequence similarity homeolog pairs. A *k* – nearest neighbor classification (knn) approach was then applied to the dataset using this training set. The *k* - nearest neighbor method uses votes from the training set to classify, therefore it supplies an objective method not only to place protokaryotypes into either a tetrasomic or disomic class, but also to identify intermediates and towards which class they are more similar based on the number and kind of votes received from the training set. In order to establish the *k* nearest-neighbors for each species, we used the resampling-based approach of 10-fold cross-validation repeated for 100 iterations implented in the trainControl function in the R package caret (v6.0-84, Kuhn 2019). For a *k* of one to 10, the median sequence similarity between all protokaryotypes for each species was divided into 10 folds, with the first fold used to test the model and the remaining folds to train the model. Next, the second fold was used to test the model and the other folds were the training set. This process continued for a total of 10 times and was repeated 100 times. The *k* for each species was chosen based on the largest number of *k* nearest-neighbors exhibiting the highest accuracy from the cross-validation procedure. This *k* was then used to classify homeologous pairs as disomic or tetrasomic, along with the predefined training set (*knn* function, Ripley and Venables 2019). Overall, similarity between the two predicted categories from the knn classification was tested with a Wilcoxon test (R Core Team 2018) incorporating percent similarity from all alignments to determine whether the categories displayed significantly different sequence similarity between homeologs (alpha = 0.01). All scripts used in this analysis are available on GitHub (see *Data Availability*).

## Data Availability

Raw sequence data has been uploaded to SRA under BioProject PRJNA555579. File S1 contains detailed descriptions of all supplemental files. File S2 contains sampling information for cisco (*C. artedi*) families. File S3 contains the Male linkage map for cisco (*C. artedi*). File S4 contains information for each marker on the female and male cisco (*C. artedi*) linkage maps. File S5 contains homologous chromosome arms determined by MapComp. File S6 contains the probable metacentric chromosomes from the MapComp analysis for coregonines. File S7 contains homeologous chromosome pairs for currently available haploid linkage maps. File S8 contains all the homeologous chromosome pairs for all available salmonid genomic resources. File S9 contains support for classifications from k – nearest neighbor machine learning algorithm. Code used to generate the Linkage mapping is available at https://github.com/DaniBlumstein/Cisco-Linkage-Map. Code used to collect *Coregonus* maps and running MapComp is available at https://github.com/bensutherland/coregonus_mapcomp. Code used for classifications from *k* nearest-neighbor machine learning algorithm is available at https://github.com/MacCampbell/residual-tetrasomy

## Results

### RADseq, SNP discovery, and data filtering

RADseq data were obtained from 746 cisco across seven families, with an average of 4.1 M reads per individual (range: 1.1 – 30.8 M reads per individual). Individuals that were genotyped at more than 30% of loci and loci that were genotyped in more than 75% of the total individuals were retained, resulting in a dataset of 676 individuals (n = 333 diploid offspring; 330 haploid offspring; and 13 parents) and 49,998 unique polymorphic loci (Supplementary file S2).

### Linkage Mapping

A total of 22,020 unique loci were mapped in the female (Figure 1) and male linkage maps (Supplementary file S2) and 27,978 loci were unplaced. The female map included 20,292 loci distributed across 38 LGs (Table 1), the male map included 6,340 loci distributed across 40 LGs, and 4,612 loci were present on both maps (Supplementary file S3). A total of 40 chromosomes was expected from karyotyping of coregonine fishes from the Great Lakes (Phillips *et al.* 1996), which matches the number of LGs mapped in males. However, male LGs 39 and 40 contained relatively few markers and may be fragments of other linkage groups rather than the two linkage groups that were not mapped in females. Eight LGs (i.e., Cart01 – Cart08) were identified as metacentric based on homology to two chromosome arms in other salmonids using MapComp (*see below*). In the female map, metacentric LGs were on average 85.44 cM (57.56 – 101.35 cM) and contained and average of 731.5 loci (range: 592 – 856). Putative acrocentric LGs in the female map were on average 59.10 cM (50.97 - 64.53 cM) and contained and average of 484 loci (range: 292 – 582). The total length of the female map was 2,456.51 cM. The average lengths of metacentric LGs on the male map were 66.76 cM (51.62 – 87.14 cM) and they contained 212 loci on average (range: 159 – 278). Putative acrocentric LGs in the male map were on average 57.00 cM (40.54 – 83.66 cM) and contained 145 loci (range: 41 – 224). The total length of the male map was 2,357.97 cM. We identified 3,383 putatively duplicated loci on the female linkage map, and of these, 2,671 loci mapped to one paralog and 709 loci mapped to both paralogs.

**Table 1.** Linkage map results for the female and male cisco (*Coregonus artedi*) linkage maps. Duplicated and non-duplicated loci are from the female linkage map, and % duplicated is the percentage of duplicated loci on each female linkage group (LG). LG type is denoted with acrocentric (A) and metacentric (M).

| C. art<br>LG | Duplicated<br>Loci | Non-duplicated<br>Loci | Length |  | Total |  | Loci/cM |  | LG<br>type |
| --- | --- | --- | --- | --- | --- | --- | --- | --- | --- |
|  |  |  | Female<br>(cM) | Male<br>(cM) | Female | Male | Female | Male |  |
| Cart01 | 203 | 510 | 101.35 | 87.14 | 713 | 224 | 7.03 | 2.57 | M |
| Cart02 | 380 | 384 | 97.77 | 61.11 | 764 | 159 | 7.81 | 2.60 | M |
| Cart03 | 223 | 477 | 93.56 | 78.98 | 700 | 248 | 7.48 | 3.14 | M |
| Cart04 | 45 | 547 | 92.45 | 67.80 | 592 | 194 | 6.40 | 2.86 | M |
| Cart05 | 272 | 584 | 91.73 | 58.07 | 856 | 219 | 9.33 | 3.77 | M |
| Cart06 | 104 | 729 | 91.36 | 68.95 | 833 | 278 | 9.12 | 4.03 | M |
| Cart07 | 184 | 496 | 57.72 | 60.40 | 680 | 204 | 11.78 | 3.38 | M |
| Cart08 | 182 | 532 | 57.56 | 51.62 | 714 | 170 | 12.40 | 3.29 | M |
| Cart09 | 230 | 271 | 64.53 | 44.47 | 501 | 92 | 7.76 | 2.07 | A |
| Cart10 | 18 | 488 | 63.98 | 49.68 | 506 | 152 | 7.91 | 3.06 | A |
| Cart11 | 36 | 510 | 63.72 | 49.41 | 546 | 165 | 8.57 | 3.34 | A |
| Cart12 | 370 | 177 | 63.18 | 74.69 | 547 | 80 | 8.66 | 1.07 | A |
| Cart13 | 24 | 558 | 62.90 | 53.67 | 582 | 181 | 9.25 | 3.37 | A |
| Cart14 | 68 | 454 | 62.66 | 59.74 | 522 | 140 | 8.33 | 2.34 | A |
| Cart15 | 87 | 318 | 62.45 | 48.26 | 405 | 121 | 6.49 | 2.51 | A |
| Cart16 | 22 | 456 | 62.25 | 45.80 | 478 | 153 | 7.68 | 3.34 | A |
| Cart17 | 27 | 478 | 62.25 | 53.19 | 505 | 160 | 8.11 | 3.01 | A |
| Cart18 | 22 | 446 | 61.68 | 42.08 | 468 | 136 | 7.59 | 3.23 | A |
| Cart19 | 27 | 541 | 61.85 | 75.08 | 568 | 213 | 9.18 | 2.84 | A |
| Cart20 | 28 | 536 | 60.19 | 48.36 | 564 | 224 | 9.37 | 4.63 | A |
| Cart21 | 30 | 388 | 59.04 | 61.23 | 418 | 142 | 7.08 | 2.32 | A |
| Cart22 | 19 | 544 | 58.96 | 49.49 | 563 | 204 | 9.55 | 4.12 | A |
| Cart23 | 49 | 473 | 58.57 | 55.39 | 522 | 162 | 8.91 | 2.92 | A |
| Cart24 | 11 | 556 | 58.38 | 57.23 | 567 | 195 | 9.71 | 3.41 | A |
| Cart25 | 23 | 458 | 58.28 | 51.08 | 481 | 161 | 8.25 | 3.15 | A |
| Cart26 | 28 | 445 | 58.09 | 50.91 | 473 | 147 | 8.14 | 2.89 | A |
| Cart27 | 30 | 499 | 58.00 | 60.69 | 529 | 213 | 9.12 | 3.51 | A |
| Cart28 | 24 | 444 | 57.79 | 54.44 | 468 | 141 | 8.10 | 2.59 | A |
| Cart29 | 13 | 404 | 57.78 | 62.64 | 417 | 149 | 7.22 | 2.38 | A |
| Cart30 | 27 | 384 | 57.76 | 71.50 | 411 | 141 | 7.12 | 1.97 | A |
| Cart31 | 22 | 541 | 56.93 | 77.82 | 563 | 184 | 9.89 | 2.36 | A |
| Cart32 | 15 | 478 | 56.56 | 75.17 | 493 | 177 | 8.72 | 2.35 | A |
| Cart33 | 19 | 460 | 56.37 | 49.57 | 479 | 146 | 8.50 | 2.95 | A |
| Cart34 | 260 | 82 | 55.54 | 83.66 | 342 | 62 | 6.16 | 0.74 | A |
| Cart35 | 38 | 254 | 54.74 | 40.54 | 292 | 81 | 5.33 | 2.00 | A |
| Cart36 | 18 | 332 | 54.45 | 45.50 | 350 | 117 | 6.43 | 2.57 | A |
| Cart37 | 23 | 402 | 53.16 | 55.82 | 425 | 122 | 7.99 | 2.19 | A |
| Cart38 | 24 | 431 | 50.97 | 73.21 | 455 | 171 | 8.93 | 2.34 | A |
| Cart39 | 0 | 0 | 0.00 | 45.13 | 0 | 41 | 0.00 | 0.91 | A |
| Cart40 | 0 | 0 | 0.00 | 58.46 | 0 | 71 | 0.00 | 1.21 | A |
| Average | 80.63 | 426.68 | 61.41 | 58.95 | 507.30 | 158.50 | 7.89 | 2.73 |  |
| Total | 3225 | 17067 | 2456.51 | 2357.98 | 20292 | 6340 | 315.42 | 109.34 |  |

**Figure 1.**
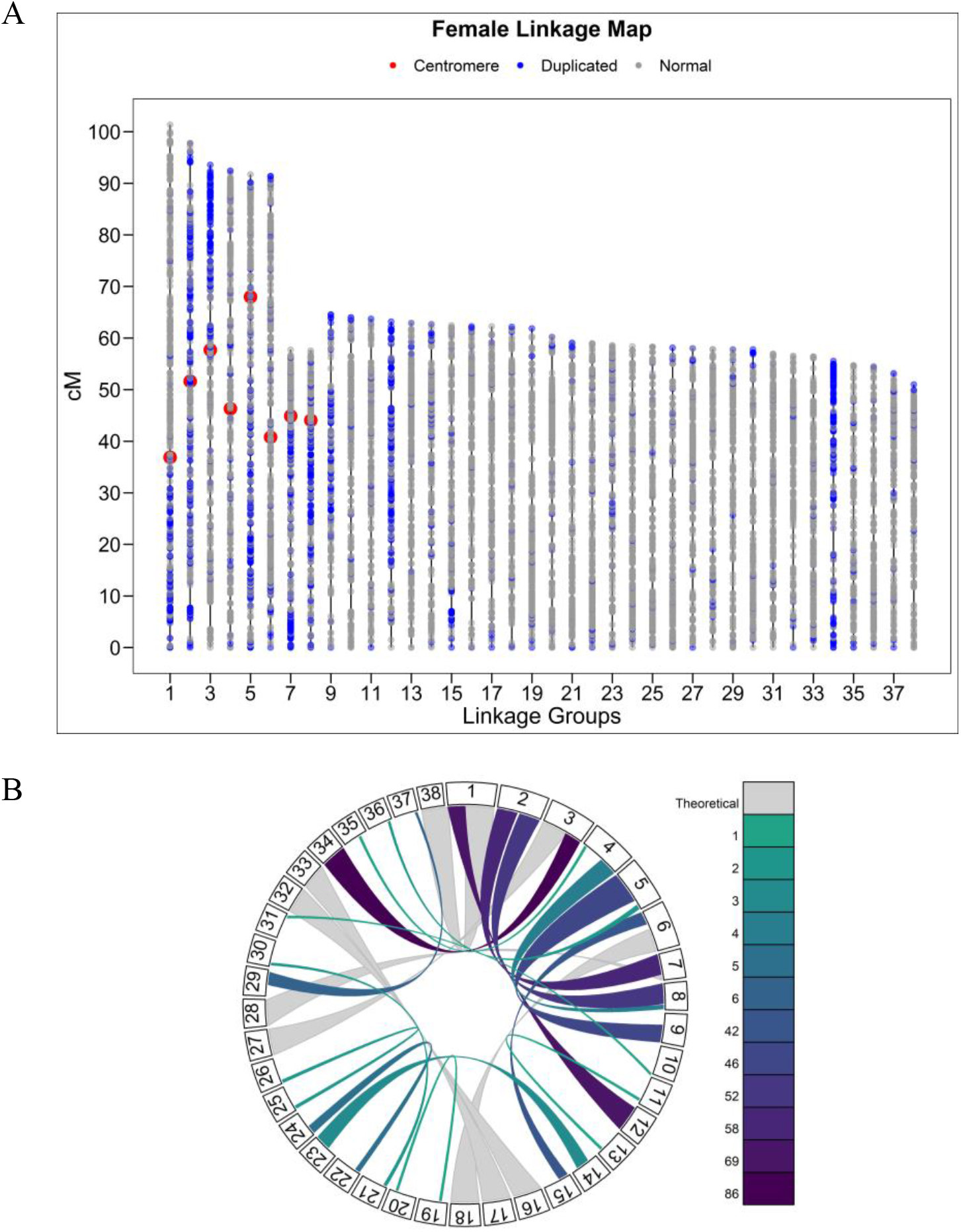
A) Female linkage map for cisco (*Coregonus artedi*) containing 20,929 loci. Each dot represents a locus, duplicated loci are blue and non-duplicated loci are gray. Lengths are in centimorgans (cM). Approximate location of centromeres for metacentric LGs are denoted in red. Metacentric LGs were identified through homologous relationships of chromosome arms with other Salmonids via MAPCOMP. B) Circos plot of cisco LGs highlighting 17 supported homeologous regions within the linkage map. Included in the 17 homeologous regions are six of the eight regions that are likely still residually tetrasomic across the Salmonids. Colors represent the number of markers supporting relationship, with darker colors representing higher marker numbers (maximum support = 86 markers) and theoretical links inferred via MAPCOMP.

### Comparative analysis of syntenic regions of linkage maps via MapComp

The main focus of our comparative analysis was to define the homologous and homeologous relationships among the linkage groups available for the coregonines, specifically in cisco (current study), lake whitefish (Gagnaire *et al.* 2013), and European whitefish (De-Kayne and Feulner 2018), and bring these species into the context of the broader chromosomal correspondence within the lineage by identifying the homologous chromosome arms in brook trout, Atlantic salmon, and Chinook salmon, as well as the non-duplicated northern pike (Table 2, Supplementary file S5). To facilitate these comparisons, we applied the same chromosome identification system as used by Sutherland *et al.* (2016), here termed the “protokaryotype identifier (PK)” system. For consistency, we maintain the .1 and .2 definitions for each ancestral chromosome pair (PK) as used in the prior work (Sutherland *et al.* 2016).

**Table 2.** MAPCOMP results documenting homologous chromosomes for the three coregonines; cisco (C. art), lake whitefish (C. clu, Gagnaire *et al.* 2013), and European whitefish (C. alb, De-Kayne and Feulner 2018), integrated with Atlantic salmon (S. sal, Lien *et al.* 2016), brook trout (S. fon, Sutherland *et al.* 2016), and Chinook salmon (O. tsh, Brieuc *et al.* 2014). Homologous chromosomes for all species are named using the corresponding Northern Pike (E. luc) linkage group as a reference (Rondeau *et al.* 2014), as per Sutherland *et al.* (2016), here termed protokaryotype ID (PK). Letters after linkage group (LG) names indicate the first (a) or second (b) arm of the LG, ^ indicates weak evidence and * indicates uncertainty between homeologs from MAPCOMP analysis.

| **E. luc**<br>**(PK)** | **C. art** | **C. alb** | **C. clu** | **S. fon** | **S. sal** | **O. tsh** |
| --- | --- | --- | --- | --- | --- | --- |
| **01.1** | Cart23 | Calb16 | Cclu28 | Sf25 | Ssa20b | Ots13q |
| **01.2** | Cart14 | Calb33 | Cclu35 | Sf38 | Ssa09c | Ots14q |
| **02.1** | Cart01a | Calb02b^* | Cclu04a | Sf06a | Ssa26 | Ots04q |
| **02.2** | Cart12 | Calb02b^* | Cclu04a^* | Sf28 | Ssa11a | Ots12q |
| **03.1** | Cart25 | Calb19 | Cclu25 | Sf22 | Ssa14a | Ots10q |
| **03.2** | Cart26 | Calb22 | Cclu26 | Sf11 | Ssa03a | Ots28 |
| **04.1** | Cart30 | Calb29 | Cclu16 | Sf33 | Ssa09b | Ots08q |
| **04.2** | Cart21 | Calb30 | Cclu29 | Sf07b | Ssa05a | Ots21 |
| **05.1** | Cart06b | Calb01a | Cclu05a | Sf01a | Ssa19b | Ots24 |
| **05.2** | Cart18 | Calb35 or Calb40 | Cclu15 | Sf27 | Ssa28 | Ots25 |
| **06.1** | Cart06a | Calb01b | Cclu05b | Sf01b | Ssa01b | Ots01q |
| **06.2** | Cart15 | Calb27 | Cclu05b^* | Sf36 | Ssa18a | Ots06q |
| **07.1** | Cart20 | Calb06 | Cclu13 | Sf08b | Ssa13b | Ots09p |
| **07.2** | Cart19 | Calb07 | Cclu08 | Sf09 | Ssa04b | Ots30 |
| **08.1** | Cart27 | Calb17 | Cclu36 | Sf04a | Ssa23 | Ots01p |
| **08.2** | Cart03a | Calb08 | Cclu06a | Sf17 | Ssa10a | Ots05q |
| **09.1** | Cart03b* | not identified | Cclu06b | Sf42 | Ssa02b | Ots32 |
| **09.2** | Cart34* | not identified | not identified | Sf03b | Ssa12a | Ots02q |
| **10.1** | Cart08a | Calb20b | Cclu10 | Sf23 | Ssa27 | Ots13p |
| **10.2** | Cart04b | Calb09a | Cclu24a | Sf34 | Ssa14b | Ots31 |
| **11.1** | Cart05a | Calb13b | Cclu18 | Sf14 | Ssa06a | Ots27 |
| **11.2** | Cart09 | Calb34 | not identified | Sf08a | Ssa03b | Ots09q |
| **12.1** | Cart33 | Calb14 | Cclu27 | Sf18 | Ssa13a | Ots22 |
| **12.2** | Cart16 | Calb28 | Cclu14 | Sf30 | Ssa15b | Ots16q |
| **13.1** | Cart17 | Calb25 | Cclu34 | Sf06b | Ssa24 | Ots04p |
| **13.2** | Cart32 | Calb31 | Cclu37 | Sf40 | Ssa20a | Ots12p |
| **14.1** | Cart01b | Calb02a | Cclu04b | Sf13 | Ssa01c | Ots20 |
| **14.2** | Cart38 | Calb11 | Cclu33 | Sf10 | Ssa11b | Ots33 |
| **15.1** | Cart10 | Calb18 | Cclu31 | Sf35 | Ssa09a | Ots08p |
| **15.2** | Cart31 | Calb10 | Cclu22 | Sf12 | Ssa01a | Ots11q |
| **16.1** | Cart07a | Calb03b | Cclu02b or Cclu03 | Sf26 | Ssa21 | Ots26 |
| **16.2** | Cart28 | Calb21 | Cclu32 | Sf24 | Ssa25 | Ots03q |
| **17.1** | Cart24 | Calb05 | Cclu38 | Sf03a | Ssa12b | Ots02p |
| **17.2** | Cart22 | Calb12 | Cclu21 | Sf21 | Ssa22 | Ots07q |
| **18.1** | Cart29 | Calb24 | Cclu40 | Sf19 | Ssa15a | Ots05p |
| **18.2** | Cart37 | Calb23 | Cclu17 | Sf31 | Ssa06b | Ots18 |
| **19.1** | Cart13 | Calb04 | Cclu30 | Sf15 | Ssa10b | Ots19 |
| **19.2** | Cart11 | Calb15b | Cclu11 | Sf20 | Ssa16a | Ots06p |
| **20.1** | Cart08b^ | Calb36 | Cclu01a | Sf07a | Ssa05b | Ots23 |
| **20.2** | Cart02b | Calb20a | not identified | Sf29 | Ssa02a | Ots03p |
| **21.1** | Cart05b | Calb13a | Cclu12 | Sf05b | Ssa29 | Ots29 |
| **21.2** | Cart36 | Calb26 | Cclu39 | Sf16 | Ssa19a | Ots16p |
| **22.1** | Cart02a | Calb39^ | not identified | Sf39 | Ssa17a | Ots07p |
| **22.2** | not identified | Calb15a | Cclu19^ | Sf05a | Ssa16b | Ots14p |
| **23.1** | Cart07b^* | Calb03a | Cclu02a | Sf02b | Ssa07b | Ots15p |
| **23.2** | Cart07b^* | missing | Cclu01b^ | Sf37 | Ssa17b | Ots17 |
| **24.1** | Cart04a | Calb09b | Cclu24b | Sf02a | Ssa07a | Ots15q |
| **24.2** | Cart35 | Calb32 | Cclu23 | Sf32 | Ssa18b | Ots10p |
| **25.1** | not identified | not identified | Cclu09^ | Sf04b | Ssa04a | Ots34 |
| **25.2** | not identified | not identified | not identified | Sf41 | Ssa08a | Ots11p |

In brief, PKs correspond to hypothetical ancestral salmonid chromosomes, which are thought to be similar to the salmonid WGD sister outgroup, the Esociformes (Ishiguro *et al.* 2003, López *et al.* 2004) and are ordered as PK 01-25. Protokaryotypes correspond 1:1 with the northern pike genome but have two descendant homeologous regions within salmonid genomes. For example, PK 01 corresponds to northern pike chromosome 01 and was an ancestral pre-duplication salmonid chromosome which gave rise to homeologous Atlantic salmon chromosomes Ssa09c (PK 01.2) and Ssa20b (PK 01.1) and to homeologous rainbow trout Omy27 (PK 01.1) and Omy24 (PK 01.2) (Supplementary file S8; Sutherland *et al.* 2016). PKs in the previously hypothesized “magic eight” PKs from linkage mapping studies are PKs 02, 06, 09, 11, 20, 22, 23, 25. PKs defined as LORes by Robertson *et al.* (2017) and those that displayed residual tetraploidy in previous genome-based studies (Lien *et al.* 2016; Campbell *et al.* 2019) are the same as these with the exception of PK 06, which is not identified as residually tetraploid.

Most PKs were identifiable in the MapComp analysis conducted here, with some notable exceptions for each coregonine species. In cisco, chromosome arms PK 22.2, 25.1 and 25.2 were unidentified; two of these arms (PK 25.1 and 25.2) were also unidentified in the European whitefish linkage map (De-Kayne and Feulner 2018). Additionally, it was difficult to determine correspondences for PK 09 and 23. In European whitefish, five chromosome arms were unidentifiable (PK 09.1, 09.2, 23.2, 25.1 and 25.2), and there were homeology ambiguities for PK 02, as well as homology ambiguities for PK 05.2 (Table 2). In lake whitefish, five arms were unidentifiable (i.e., PK 09.2, 11.2, 20.2, 22.1, and 25.2), and there were homeology ambiguities for PK 02.2 and 06.2. In multiple species, arms where it was difficult to determine homologous relationships often had a high proportion of duplicated loci, presumably making distinguishing homologs and homeologs challenging. Nonetheless, most homologs and homeologs (42/50; 84%) were identified in all three coregonine species. This information was then leveraged to characterize the fusion/fission history within the Coregoninae lineage using the methods outlined in Sutherland *et al.* (2016).

The fusion/fission analysis indicated far fewer species-specific fusions than identified for subfamily Salmoninae in Sutherland *et al.* (2016), with most fusions that occurred within subfamily Coregoninae occurring prior to the divergence of the coregonines (Figure 2). This difference in species-specific fusions may also be related to the general lower number of fusions in coregonines relative to Salmo and Oncorhynchus (Supplementary file S6); although, in the coregonines the majority of fusions were observed in more than one species, which was not observed in most other species previously characterized. Two strongly supported fusions were observed in all three coregonine species: fusions PK 05.1-06.1 and 10.2-24.1. PK 11.1-21.1 was fused in both cisco and European whitefish, which presumably underwent a fission in lake whitefish (Figure 2). However, evidence for the correspondence for lake whitefish for PK 11.1 and 21.1 was not highly conclusive, and a more recent analyses of lake whitefish by regenerating the linkage map suggests that PK 11.1-21.1 may have not underwent a fission in this species and is indeed still fused (Claire Mérot, pers. comm.). Therefore, more work is needed to determine whether this fusion is conserved in all three species. The full characterization of fissions will require the resolution of the ambiguous arms that are considered as probable in the current analysis, and this may be further clarified in future work.

**Figure 2.**
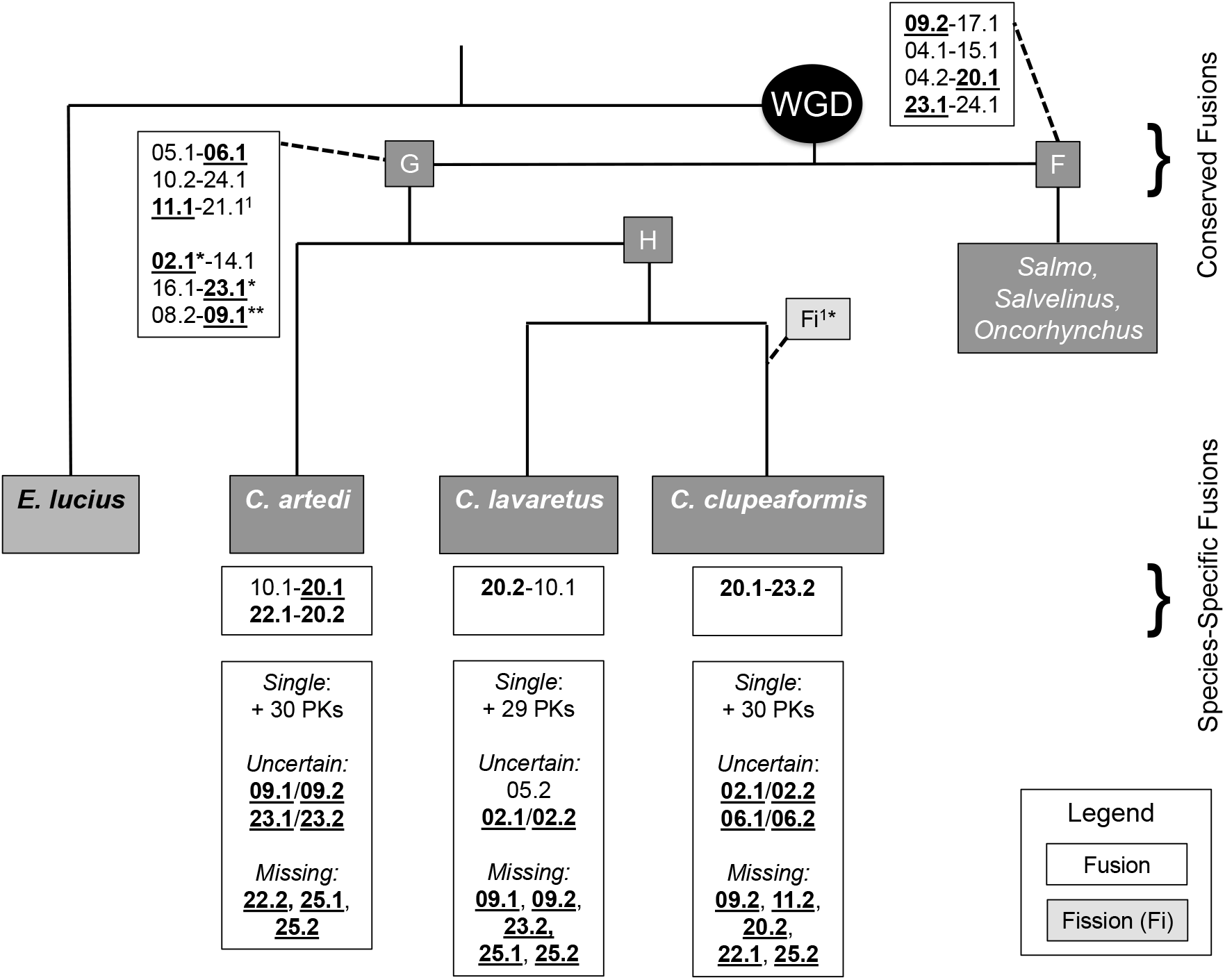
Fusions and fissions in the Coregoninae and Salmoninae lineages. This is an extension of Figure 4 from Sutherland *et al.* (2016). White boxes display the fusion events, where the homologous chromosomes for all species are named according to the protokaryotype ID. Bold and underlined chromosome numbers are the homeologous pairs that exhibit residual tetraploidy (i.e., “magic eight”), * indicate uncertainty in one species, and ** indicates uncertain in two species (i.e., *C. artedi* is ambiguous for homeolog 09.1 or 09.2 while *C. lavaretus* is missing 09.1 and 09.2). Above the species names are conserved fusions, whereas below are the species-specific fusions. The phylogeny is adapted from (Crete-Lafreniere *et al.* 2012). Branch lengths do not represent phylogenetic distance, only relative phylogenetic position. 1Arms 11.1-21.1 were fused in both *C. artedi* and *C. lavaretus*, but likely underwent fission in lake whitefish (but see Results).

In summary, five fusions were likely shared among all three species, and one was shared between cisco and European whitefish, and possibly all three species (PK 11.1-21.1). Cisco had two species-specific fusions (PK 10.1-20.1 and 22.1-20.2), bringing the total count of observed fusions to eight. European whitefish had one species-specific fusion (PK 20.2-10.1), bringing the total count of observed fusions to seven. Lake whitefish also had one species-specific fusion (PK 20.1-23.2), bringing the total count to six. Interestingly, the PK 09.2-17.1 fusion that was originally proposed to be shared among all known salmonids (Sutherland *et al.* 2016), was found not to be fused in any of the species here, suggesting either that this fusion occurred after the divergence of *Coregonus* from the ancestor of the rest of the salmonids, or that a fission occurred at the base of the coregonines (Figure 2). The observation that this fusion was not present in grayling Varadharajan *et al.* 2018) suggests the former.

### Homeolog identification, similarity, and inheritance mode

A second major goal of this study was to compare homeologous relationships and modes of inheritance within and among species. We identified 17 of the 25 homeologous chromosome pairs (PK) in cisco using the markers that could be mapped to both homeologs in the linkage map, and each homeologous pair shared between one and 86 duplicated loci (Fig. 3, Supplementary file S7). Of the 17 homeolog pairs, six (PK 02, 06, 09, 11, 20, and 23) had many loci (42-86) supporting homeology; these are six of the “magic eight” discussed above. The other 11 had few markers supporting homeology (i.e., 1-6) and are not members of the “magic eight”. The other two arms found in the “magic eight” were not identifiable in cisco. All of the previously constructed linkage maps for salmonids that included duplicated regions had a large number of markers supporting homeology for the “magic eight” with the exception of pink salmon, where seven of the eight PKs had high support (34 – 68 loci) but one pair (PK25) displayed substantially lower support (nine loci) (Tarpey *et al.* 2017) (Figure 3, Supplementary file S7).

**Figure 3.**
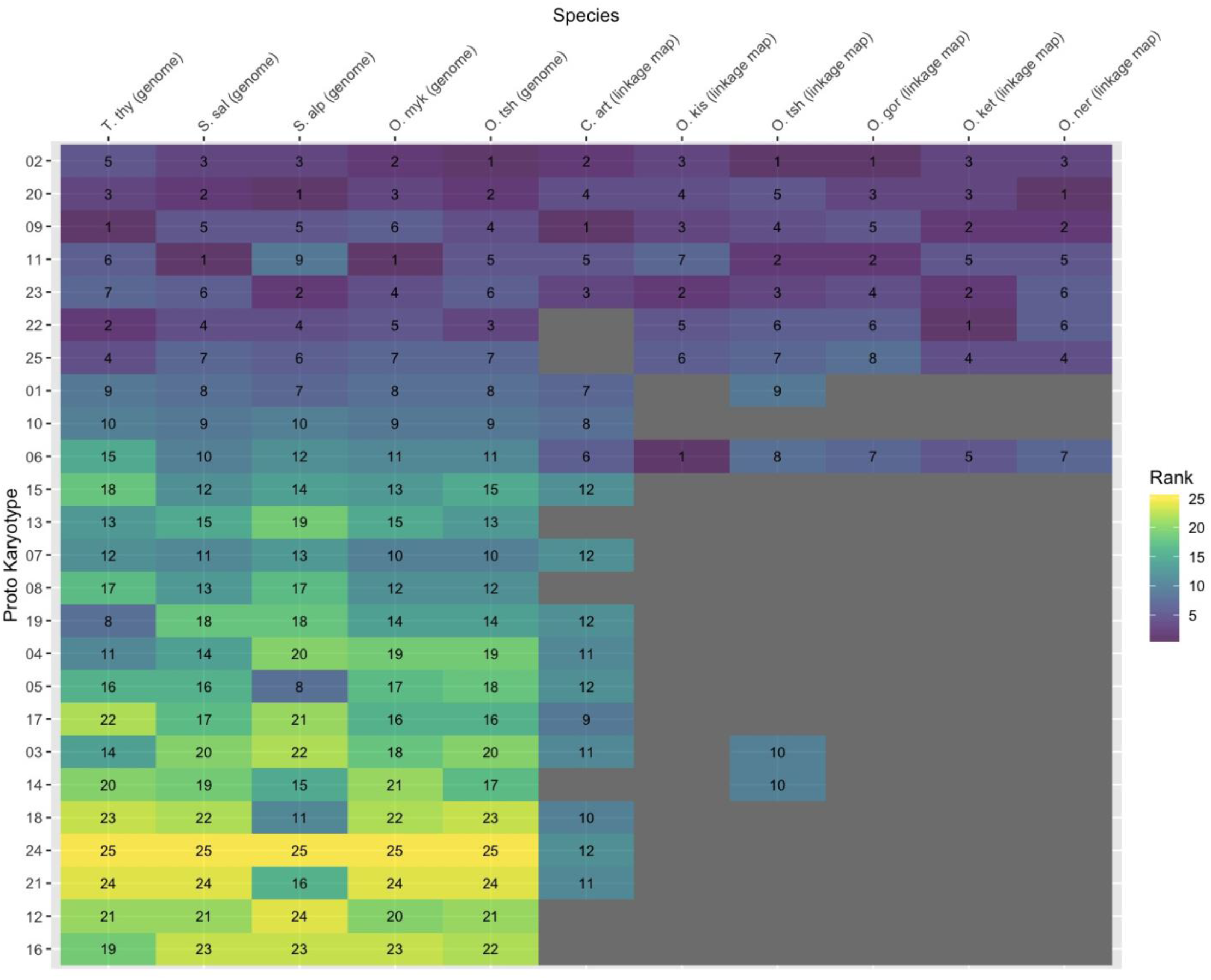
Ranking of homeologous chromosome pairs based on putative residual tetrasomic inheritance as measured by the number of markers shared among homeologs for linkage maps or percent sequence similarity for genomes. A lower rank represents more marker pairs supporting a homeolog and or a higher sequence similarity. Chromosomes for all species are named according to the protokaryotype ID (PK). PKs are ordered in the figure by averaging the ranks across all species and then sorting the averages from smallest to largest (i.e., ordered from highest support for residual tetrasomy to lowest). Grey indicates that no duplicated loci could be mapped to both homeologs. Species abbreviations are grayling (T. thy), Atlantic salmon (S. sal), Arctic char (S. alp), rainbow trout (O. myk), Chinook salmon (O. tsh), cisco (C. art), and coho salmon (O. kis).

To better understand the genetic similarity between homeologs and infer inheritance mechanisms (i.e., residual tetrasomy or disomy), all 25 known homeologous relationships were compared in reference genomes for grayling, Atlantic salmon, Arctic char, rainbow trout, and Chinook salmon (Figures 3 and 4, Supplementary file S8). Using the machine learning algorithm (see Methods), the optimal *k* nearest-neighbor for each species was identified as five. Those five nearest neighbors from the training sets voted on the assignment of a particular PK to either putatively tetrasomic or disomic classes (Figure 4), and the proportion of votes supporting each assignment are reported in Supplementary file S9. The highest observed vote proportion for assignment to a class is 4 of 5 as a result of the limit on training set size to four of each class and the five optimal *k* nearest-neighbors indicated for accuracy.

**Figure 4.**
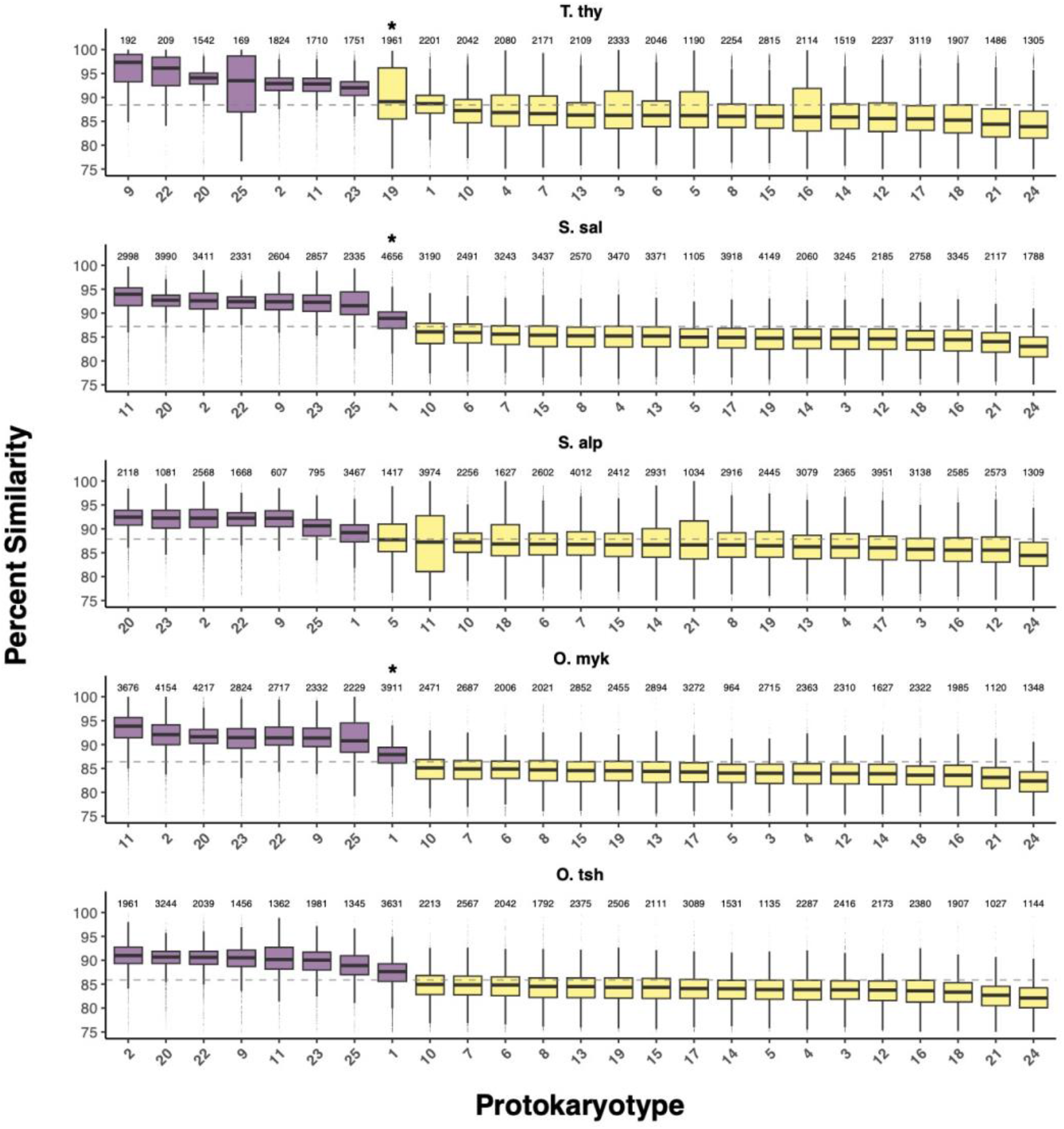
Distribution of protokaryotype (PK) similarity in aligned sections between the homeolog pairs across salmonids based on genome assemblies. For each species with a genome sequence, the percent similarity (*y* – axis) of the 25 PK pairs as shown as box plots. PK pairs are ranked from highest to lowest median similarity for each species (*x* – axis), with the average similarity of protokaryotypes presented as a dashed line. The classification of protokaryotypes by the machine learning approach described in the main text into putatively tetrasomic and disomic pairs is shown through coloring of the boxplots into purple (putatively tetrasomic) and yellow (putatively disomic). The number of alignments used in computing similarity is presented at the top of each bar. Those protokaryotypes that did not receive the highest observed voting proportion for the assigned class are indicated with an asterisk (*). PKs with high variance (e.g. PK 11 in S. alp) may be due to methodological limitations that have caused additional non-homeologous chromosome arms to be included in the comparisons (see discussion). Species abbreviations are grayling (T. thy), Atlantic salmon (S. sal), Arctic char (S. alp), rainbow trout (O. myk), and Chinook salmon (O. tsh)

For Atlantic salmon and all *Oncorhynchus* spp. (i.e., rainbow trout and Chinook salmon), the same eight PKs (i.e., PK 01, 02, 09, 11, 20, 22, 23, 25) were classified as tetrasomic using the machine learning approach. This list of PKs includes all of those defined as LORes by Robertson *et al.* (2017), and one additional (i.e., PK 01), but does not include PK 06, which is considered to be part of the “magic eight” using linkage map evidence. Arctic char showed evidence for residual tetrasomy in seven of these eight PKs, with the exception of PK 11 (see below for details regarding this discrepancy due to other chromosome arms in this fusion). Grayling also shared seven of the eight residually tetraploid homeolog pairs, with the exception of PK 01. Most PKs received the highest possible vote proportions for their classifications (0.8), however, PK01 in Atlantic salmon and rainbow trout demonstrated a lower vote proportion (0.6) (Figure 4, Supplementary file S9), suggesting reduced support (i.e., lower sequence similarity) for this homeologous pair being tetrasomic. Additionally, PK 19 in grayling, did not have the highest vote proportion and was assigned as diploid but had the highest sequence similarity in that class (Figure 4, Supplementary file S9). Sequence similarity was significantly higher for the tetrasomic PKs across all species (P < 0.0001).

Although the group of tetrasomic PKs was largely conserved across species, there was substantial variation in the relative sequence similarity between these homeolog pairs (i.e., order of highest to lowest similarity) among species. PK 01 consistently displayed the lowest sequence similarity of all the PKs in all five species where it was classified as tetrasomic and did not always receive the highest observed vote proportion (see above). However, there were a number of other homeolog pairs that displayed highly variable sequence similarity rankings across species (Figure 4). For example, PK 09 had the highest sequence similarity in the grayling genome, the sixth highest in the rainbow trout genome, and the fourth or fifth highest in the other genomes. This variation suggests that the frequency of tetravalent meiosis for each PK may differ across species and that the process of diploidization has occurred in a species-specific manner post WGD as suggested in the mechanisms proposed by Robertson *et al.* (2017).

## Discussion

The amount of genomic resources available for Salmonidae has increased drastically over the last decade. However, many previous studies investigating genome evolution in salmonids focus on one or a few species, with a limited number of studies considering broader subsets of available taxa to understand patterns of genome evolution across the Salmonidae family (but see Sutherland *et al.* 2016; Robertson *et al.* 2017). Here, we utilize genomic resources along with a newly generated high-density linkage map for cisco to compare patterns of homology, fusion/fission events, homeology, and residual tetrasomy across species. The cisco linkage map incorporates duplicated regions and contains 20,450 loci, making it denser than most salmonid RAD-based haploid linkage (typically built from 3,000 to 7,000 loci). The higher density linkage map was achieved by using an updated RAD library preparation and linkage map algorithms (Rastas 2017) in addition to including more families and more individuals per family. Higher marker density allowed the identification of orthologous relationships between coregonines and other salmonids as well as to identify homeologous chromosomes in cisco. We also demonstrate the use of the protokaryotype ID (PK), defined here but first used in Sutherland *et al.* (2016), for comparative analyses in salmonids in order to unify and facilitate comparative approaches in salmonid linkage maps and chromosome-level assemblies. Comparisons across Salmonidae revealed that patterns of rediploidization are relatively similar across genera and loosely correspond with phylogeny. However, we did identify substantial variation in sequence similarity between homeologs both within species across homeolog pairs, and among species, suggesting that frequently used binary classifications such as AORe/LORe and “magic eight” may be oversimplified.

### Protokaryotype identifiers to facilitate comparative genomics in salmonids and other fishes

Comparative genomics within Salmonidae is important for the interpretation of the effects of rediploidization after WGD on genome evolution (e.g., Berthelot *et al.* 2014; Kodama *et al.* 2014; Lien *et al.* 2016). However, chromosomes in all species have been named differently, making it difficult to directly compare studies without complicated lookup tables or alignments to confirm homology (e.g. Brieuc *et al.* 2014; Kodama *et al.* 2014). Recently developed methods for connecting linkage maps through reference genomes (Sutherland *et al.* 2016) facilitated description of homologous relationships for all linkage group arms across salmonids (with a few exceptions in coregonines). Additionally, Sutherland *et al.* (2016) and Savilammi *et al.* (2019) have explored the utility of naming chromosomes based on homology to northern pike. This naming system has the potential to facilitate comparative genomics in salmonids by creating a “Mueller element”-like system (reviewed in Schaeffer 2018), where each chromosome arm has a universal identifier. However, there also remains value in species-specific identifiers; for example, Cart03 is the third named linkage group in the Cisco linkage map (Table 2). By comparison, Cart03 named via the PK system could be Cart03 (PK 08.2-09.1) or Cart03_PK08.2-09.1_ as Cart03 represents the fusion of two ancestral salmonid chromosome arms 08.2 and 09.1 (Figure 1, Table 2).

While Sutherland *et al.* (2016) named salmonid chromosomes based on ancestral northern pike chromosomes, the utility of the system was not yet fully explored or discussed. Here, we demonstrate the utility of this system and advocate its use in future studies. For example, the PK system can facilitate comparisons of chromosomes containing genes for adaptive potential in sockeye salmon (So13PK18.2, TULP4, Larson *et al.* 2017), for run timing in Chinook (Ots28_PK03.2_, GREB1L, Prince *et al.* 2017), and for age-at-maturity in Atlantic salmon (Ssa25_PK16.2_, VGLL3, Barson *et al.* 2015). While there may be some sections of the PK that are not always retained (e.g., some transposition of parts of chromosomes), as long as the majority of the chromosome is preserved, then the PK system enables general comparisons. The PK system will facilitate quick and accurate comparisons across taxa, adding significant value to the myriad studies searching for adaptively important genes and regions in salmonids by leveraging comparative approaches. This system was previously applied by Sutherland *et al.* (2017) to compare sex chromosomes across the species by comparing chromosomes containing the transposing salmonid sex determining gene (sdY, Yano *et al.* 2012). This comparison demonstrated that some chromosome arms more frequently contain or are fused to the chromosome that contains the sex determining gene than would be expected by chance or explainable by phylogenetic conservation (i.e., PK 01.2 (AC04q), PK 03.1 (Cclu25, Co30, So09), PK 19.1 (So09.5, AC04q.1), PK 15.1 (AC04q.2, BC35), Sutherland *et al.* 2017). Even more intriguing is that the northern pike naming was based on the three-spined stickleback *Gasterosteus aculeatus* (Rondeau *et al.* 2014), and PK 19 is the sex determining chromosome in three-spined stickleback (Peichel *et al.* 2004). As observed above, this chromosome is often fused with sex chromosomes in salmon (Sutherland *et al.* 2017). By comparison, using the naming system, it is easy to observe that LG24 in northern pike (i.e., PK 24 in salmonids), recently identified to hold the sex determining gene in northern pike (Pan *et al.* 2019), does not appear to contain the sex-determining locus in any tested salmonid. Deriving this information would be more difficult without the PK system and would require extensive cross-referencing.

The example of comparing sex chromosomes from the PK system indicates a broad phylogenetic utility of this nomenclature as it applies to three-spined stickleback (a neoteleost) as well as Esociformes and Salmoniformes. The protokaryotypes as defined here may be able to represent the ancestral karyotype of the five major euteleost lineages and be applicable in comparative genomic studies among and within (1) Esociformes and Salmoniformes, (2) Stomiatii, (3) Argentiniformes, (4) Galaxiiformes, and (5) Neoteleostei (Betancur-R *et al.* 2013). Exploration of the PK system as defined here and its applicability across euteleosts should be conducted to determine the suitability of the PK system for comparative genomics in the Euteleostei.

### Homology and fission/fusion history in coregonines

Comparisons using linkage maps for three coregonine species (i.e., cisco, lake whitefish and European whitefish), allowed us to assess homology and variation in karyotypes across the genus. Within lake whitefish, aneuploidy has been documented in diverged populations and historical contingency (Dion-Côté *et al.* 2015, 2017). Our results show ambiguity in homologous relationships remained for at least five chromosome arms in all three coregonines. This degree of uncertainty was much higher than documented in *Salmo*, *Oncorhynchus*, and *Salvelinus* by Sutherland *et al.* (2016), where there were only two ambiguities across these groups. Coregonines appear to have a number of relatively small acrocentric chromosomes (Phillips and Rab 2001), some of which contain a high degree of duplicated loci, making constructing linkage maps more difficult than for other salmonids (Gagnaire *et al.* 2013; De-Kayne and Feulner 2018). For example, PK 25 has never been successfully mapped in coregonines, likely because it is small, submetacentric or acrocentric, and contains many duplicates. In an attempt to recover the missing PK in coregonines linkage maps, various approaches were attempted, including using unassigned makers to form LGs, using only non-duplicated loci from the female cisco linkage map to form LGs, and aligning unassigned sequences to reference genomes. Markers either formed very large LG with many gaps, still remained unassigned, or aligned to unplaced scaffolds on reference genomes. In other salmonids, where PK 25 is part of larger and/or metacentric chromosome, mapping is expected to be easier as there are many disomically inherited markers on the chromosome. Interestingly, the fact that PK 25 is likely residually tetrasomic in cisco, even though it is likely an acrocentric or submetacentric chromosome, indicates that metacentric chromosomes may be not required for homeologous recombination, as previously suggested in Lien *et al.* (2016). This potentially contradicts previous theory which suggests that homeologous recombination requires at least one chromosome arm to be metacentric (Kodama *et al.* 2014), but requires further testing given uncertainties regarding PK25. Additionally, PK 25.2 in grayling is a submetacentric chromosome and also displays signals of residual tetrasomy (Savilammi *et al.* 2019), potentially providing further evidence that a small secondary arm may be sufficient to facilitate tetrasomic meiosis.

The fusion history in coregonines differs substantially from many other members of the salmonid family. Members of the *Coregonus*, *Salvelinus*, and *Thymallus* genera possess the “A karyotype,” with a diploid chromosome number (2N) ~80 and many acrocentric chromosomes, whereas *Oncorhynchus* and *Salmo*, possess the “B karyotype,” with 2N ~60 and many metacentric chromosomes (Phillips and Rab 2001). Given that these both come from an ancestral type of n = 50 chromosome arms, species with the “A karyotype” have undergone fewer fusions than lineages with the “B karyotype”. Interestingly, it appears that “A karyotype” species also generally contain a lower proportion of species-specific fusions compared to “B karyotype” species, suggesting that the reduction in chromosome number and the higher frequency of metacentric chromosomes characteristic of the “B karyotype”, comes from species-specific fusions. Sutherland *et al.* (2016) investigated fusion history within many species from the *Oncorhynchus* genus and found that most species had many species-specific fusions (e.g., 17 species-specific fusions in pink salmon). However, Sutherland *et al.* (2016) only investigated one species from the *Coregonus* and *Salvelinus* genera as this was all that was available at the time of publication, and no species from *Thymallus*. Our current study is the first to investigate fusion history across multiple coregonines and illustrates that most fusions are shared among species in the *Coregonus* genus, contrasting the pattern observed in *Oncorhynchus* spp. (Sutherland *et al.* 2016). The functional effect of differing fusion histories is yet to be determined, and remains an important question differentiating species within the *Coregonus*, *Salvelinus*, and *Thymallus* genera from other salmonids. Further information from genome sequencing projects, for example the European whitefish genome (De-Kayne *et al.* 2020) should facilitate important future studies contrasting genomic processes and structure in species with differing fusion histories.

### Patterns of homeology and residual tetrasomy across salmonids

Although patterns of residual tetrasomy were generally conserved, variation within and among species was observed when examining results from linkage maps versus reference genomes. Sequence similarity analyses using reference genomes suggested that all species showed evidence for residual tetrasomic inheritance in seven homeologous pairs (PK 02, 09, 11, 20, 22, 23, and 25) with the exception of PK11 in Arctic char (see below). Using linkage maps, these same seven homeologous pairs have been found to be tetrasomic in *Oncorhynchus* (Kodama *et al.* 2014; McKinney *et al.* 2019), *Salvelinus* (Sutherland *et al.* 2016; Nugent *et al.* 2017), *Salmo* (Robertson *et al.* 2017), and likely *Coregonus* (results reported herein), strongly suggesting that tetravalent meioses can and do form between these homeologs in all investigated species to date. However, evidence for residual tetrasomy differed between linkage map and genome methods for multiple PKs, most notably PK 06, which was classified as tetrasomic in linkage mapping studies but not in genome analyses, and PK 01 which was classified as tetrasomic in genome analyses but not linkage maps. It is likely that some of these differences are the result of methodological limitations of the current approach and point to future analysis approaches that may be able to improve upon the framework presented here. This is further described below.

The observation that PK11 did not display high sequence similarity in Arctic char might suggest a difference in diploidization rates in *Salvelinus* compared to other salmonids for this homeologous pair, but it is more likely that methodological limitations prevented us from detecting residual tetrasomy, as a linkage map study in Arctic char found a high number of duplicated markers on this PK (Nugent *et al.* 2017). The percentage similarity analysis applied in the present study uses complete chromosome alignments and requires post-filtering to remove non-homeologous alignments. This method appears to be robust when chromosome arms are well defined but, PK 11 in Arctic char appears to be composed of four chromosome arms that have come together in a series of species-specific fusions (inferred from Christensen *et al.* 2018b). Since arm boundaries were not well defined, alignments in this PK produced a wide interquartile range, suggesting that, while some regions of the PK are likely undergoing residual tetrasomy, the alignments may have masked these regions by integrating over multiple chromosome arms. This would be particularly problematic if the chromosomes being compared both contained non-target chromosomes that were homeologous. To improve upon the method applied here, better definition of the breaks between chromosome fusions could be applied and this could prevent such ambiguities or noise in the sequence similarity calculated. We therefore conclude that PK 11 is likely tetrasomic in Arctic char, but that we were unable to classify it as such due to methodological limitations. The sequence similarity method applied here is generally robust, but the fusion history of the species being analyzed needs to be considered to avoid unexpected and erroneous similarity values. Ideally, only the section containing the ancestral chromosome of interest would be being compared between the homeologs. This is an avenue of method development that will be valuable for future work.

Contrastingly, the finding that PK 06 is not tetrasomic does not appear to be due to methodological limitations of our genome analysis but may be due to differences in estimating extent of residual tetraploidy between linkage mapping and genome assembly approaches. Linkage mapping in *Oncorhynchus* and *Salvelinus* consistently finds support for tetrasomic inheritance at PK 06 (Larson *et al.* 2017; Nugent *et al.* 2017), but the genome analysis conducted here and that was conducted for rainbow trout (Campbell *et al.* 2019) found that this PK displayed intermediate sequence similarity consistent with disomic homeologs. One of the ways the two approaches differ is the length of the sequence used during each analysis. The genome analysis conducted here calculated similarity by using alignments of at least 1,000 bp, whereas linkage maps compare alleles within ~100-150 bp RADtags. The short sequences analyzed by software such as *Stacks* (Rochette and Catchen 2017; Rochette *et al.* 2019) make it possible to collapse sequences into a single locus that can be mapped at both paralogs, even when sequence divergence in a given region is relatively large. This makes linkage maps a less conservative characterization method for determining residual tetrasomy. In addition, many genome assemblers applied to salmonid genomes (e.g. Chin *et al.* 2016; Koren *et al.* 2017; Ruan and Li 2019) are not optimized for paralogous regions in polyploid genomes. This could be especially problematic for genomes that combine both disomic and tetrasomic regions, such as in salmonids. The end result is that duplicated regions may be detected as single copies as a result of sequence collapse during the assembly process (Alkan *et al.* 2011; Varadharajan *et al.* 2018). If sequences do not collapse during assembly, contigs might be fragmented and misassembled in the genome, making it difficult to differentiate between homologs and homeologs (Kyriakidou *et al.* 2018). This could lead to homeologous regions being missed altogether in genome sequences, particularly in comparisons that require chromosome-level assemblies. However, the fact that support for tetrasomic inheritance in other PKs identified as tetrasomic through linkage mapping was consistent with that observed in genome analysis strongly suggests that there is something unique with PK 06 rather than a fault with the genome analyses conducted here. Perhaps, as suggested by Campbell *et al.* (2019), the PK 06 chromosome arms are returning to a diploid state faster than the other seven tetrasomic homeolog pairs or the tetrasomically inherited portion of PK 06 is smaller than other tetrasomic PKs.

Another notable difference between linkage mapping and genome analysis was the consistent classification of PK 01 as tetrasomic in the genome analysis (five of six species) but not in any linkage map. PK 01 uniformly exhibited the least similarity between tetrasomic homeologous pairs and was assigned to the putatively tetrasomic class of PKs with less certainty by the machine learning algorithm. This suggests that PK 01 may have low levels of tetrasomy. We also observed some consistent patterns of variation in sequence similarity within disomic markers. For example, homeolog pairs for PK 24 and 21 generally displayed the lowest sequence similarity, and homeolog pairs for PK 07 and 19 displayed higher similarity. Our study therefore presents additional nuances into the rediploidization process by identifying a core group of conserved tetrasomic homeologs, potentially intermediate homeologs (PK 01, 06) and consistently diverged homeologs (PK 21, 24). Future investigations can be refined to examine four well-defined categories across PKs: tetrasomic, intermediate, disomic, and most diverged. This enhanced refinement should reduce noise from the incorrect pooling of homeologs and aid in understanding the rediploidization process in salmonids.

Interestingly, more variation in sequence similarity was observed within tetrasomic homeologs than was observed in disomic homeologs. For example, PK 23 has the second highest sequence similarity in Arctic char, the fourth highest in rainbow trout, the sixth highest in Atlantic salmon, and the seventh highest in grayling. While this may be in part due to differences in genome assembly method and assembly quality, the fact that variation exists even among the highest quality genomes (Atlantic salmon and rainbow trout) suggests that rediploidization rates at tetrasomic PKs may vary among species, even though the same seven PKs are consistently classified as tetrasomic. In other words, although there appears to be a large amount of conservation of tetrasomic inheritance between species, our genome analyses also suggest some independence in the return to disomy since the three subfamilies of salmonid split ~ 50MYA.

### Conclusions

Here we provide the most complete analysis of chromosomal rearrangements in coregonines using the currently available genomic resources and a haploid linkage map for cisco. We also integrate this analysis with prior characterizations of chromosomal rearrangements in salmonids through the use of a common identifier system, the protokaryotype ID (PK), and suggest its continued use to facilitate comparative analyses of salmonids. Our study revealed that patterns of tetrasomic inheritance are largely conserved across the salmonids, but that there is substantial variation in these patterns both within and among species. For example, while the same seven PKs appear to be tetrasomically inherited across all species examined, their relative rates of sequence similarity differ within species, suggesting the potential of independent evolutionary trajectories following speciation. Additionally, we documented that analyses based on linkage maps do not identify the same tetrasomically inherited PKs as genome analyses and postulate that this may be due to inconsistencies with genome assemblies or due to differences in the length of sequence used in comparisons. This study provides important insights about the WGD in salmon and also provides a framework that can be built upon to improve our understanding of WGDs both within and beyond salmonids.

## Acknowledgements

This project was funded by the Great Lakes Restoration Initiative (GLRI). Special thanks to the USFWS crew members Chris Olds, Paul Haver, Kaley Genther, Steve Nimcheski, and Matt McLean for assistance in field sampling, USGS Great Lakes Science Center Aquatic Research Wet Lab for egg and cisco rearing, the University of Wisconsin-Stevens Point Molecular Conservation Genetics Lab for assistance in lab work, and the support of the Turing High Performance Computing cluster at Old Dominion University. Thanks to Kris Christensen, Claire Mérot, Eric Rondeau, Louis Bernatchez, two anonymous Reviewers, and the Associate Editor Andrew Whitehead for valuable and constructive comments on the manuscript. Any use of trade, product, or company name is for descriptive purposes only and does not imply endorsement by the U.S. Government.

## Supplementary tables and figures

S1. A detailed description of all supplemental files.

S2. Sampling information for cisco (*Coregonus artedi*) families collected from Lake Huron the number of individuals used. The number of offspring shown only includes those used for mapping after the removal of individuals with low sequencing coverage. The numbers of SNPs shown include those retained following quality control filtering but before inclusion in linkage mapping.

S3. Male linkage map for cisco (*Coregonus artedi*) containing 6340 loci. Each dot represents a locus. Lengths are in centimorgans (cM).

S4. Information for each marker on the female and male cisco (*Coregonus artedi*) linkage maps. Tag is the RAD tag, and marker name is the tag followed by a designation used to differentiate duplicated loci. The “Sequence P1 column” is the sequence from the single end read for each RAD tag, and the “Sequence PE” is the sequence obtained from paired-end assemblies.

S5. MapComp determination of homologous chromosome arms.

S6: Probable metacentric chromosomes from the MapComp analysis for coregonines.

S7. Homeologous chromosome pairs for currently available haploid linkage maps and the number of maker pairs supporting homeology (Kodama *et al.* 2014; Larson *et al.* 2015; Waples *et al.* 2016; Tarpey *et al.* 2017). Homologous chromosomes are named according to the corresponding Northern Pike linkage group Protokaryotype ID (PK).

S8. Homeologous chromosome pairs for all available salmonid genomic resources.

S9. Support for classifications from k – nearest neighbor machine learning algorithm. Cross validation indicated five nearest neighbors should be used for classification across species. The vote proportion for assignment to either putatively tetrasomic or putatively disomic categories are presented as the proportion of votes out of five at the top of each bar. Median percent similarity for each protokaryotype pair is presented as the *y* – axis and protokaryotype pairs are ranked from most similar to least (*x* – axis).

