## Supplementary figures and images for "Comparative genomic analyses and a novel linkage map for cisco (*Coregonus artedi*) provide insights into chromosomal evolution and rediploidization across salmonids"

### Supplemental File 3

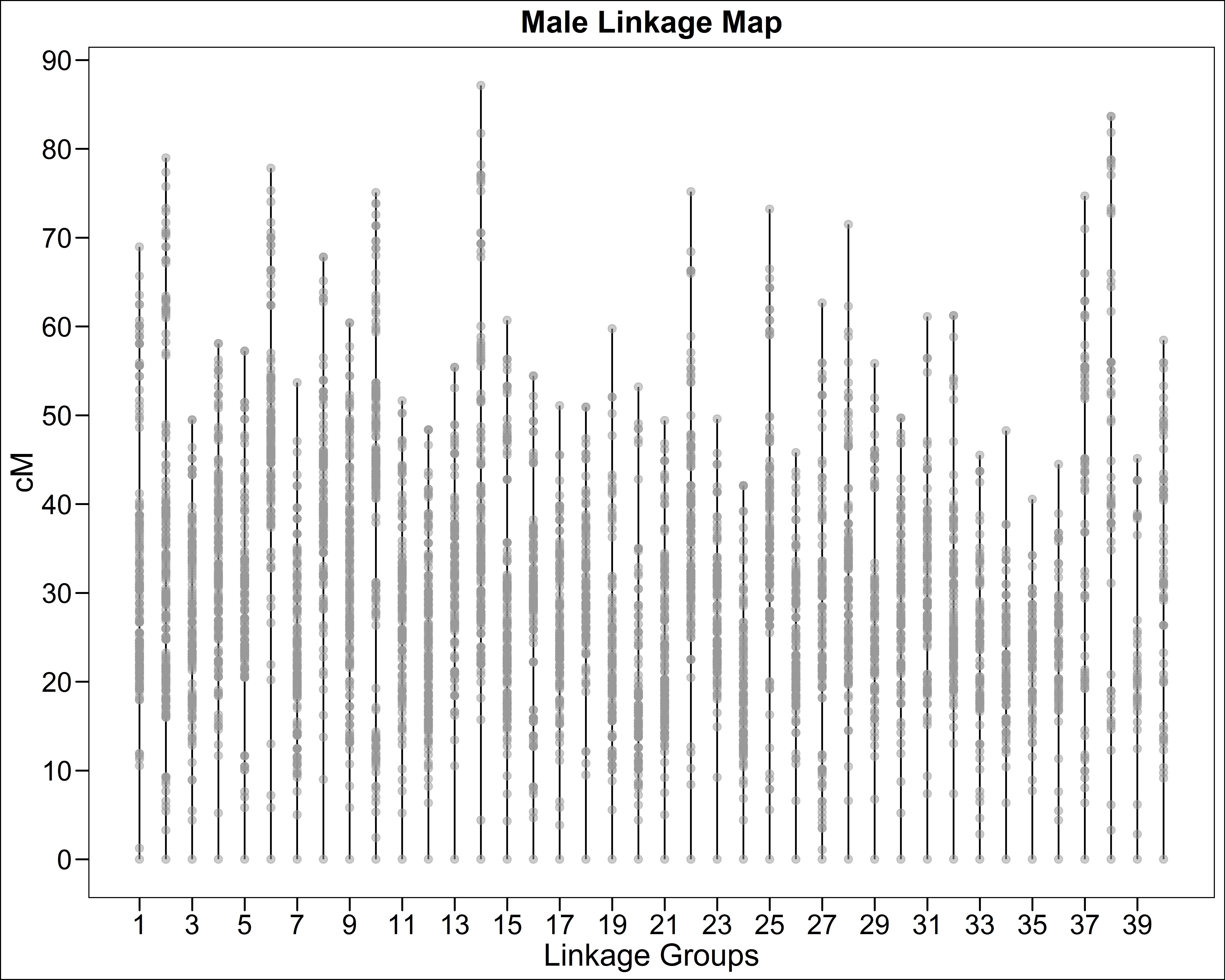

### Supplemental File 5

#### Map comparison for species Cart and Clav

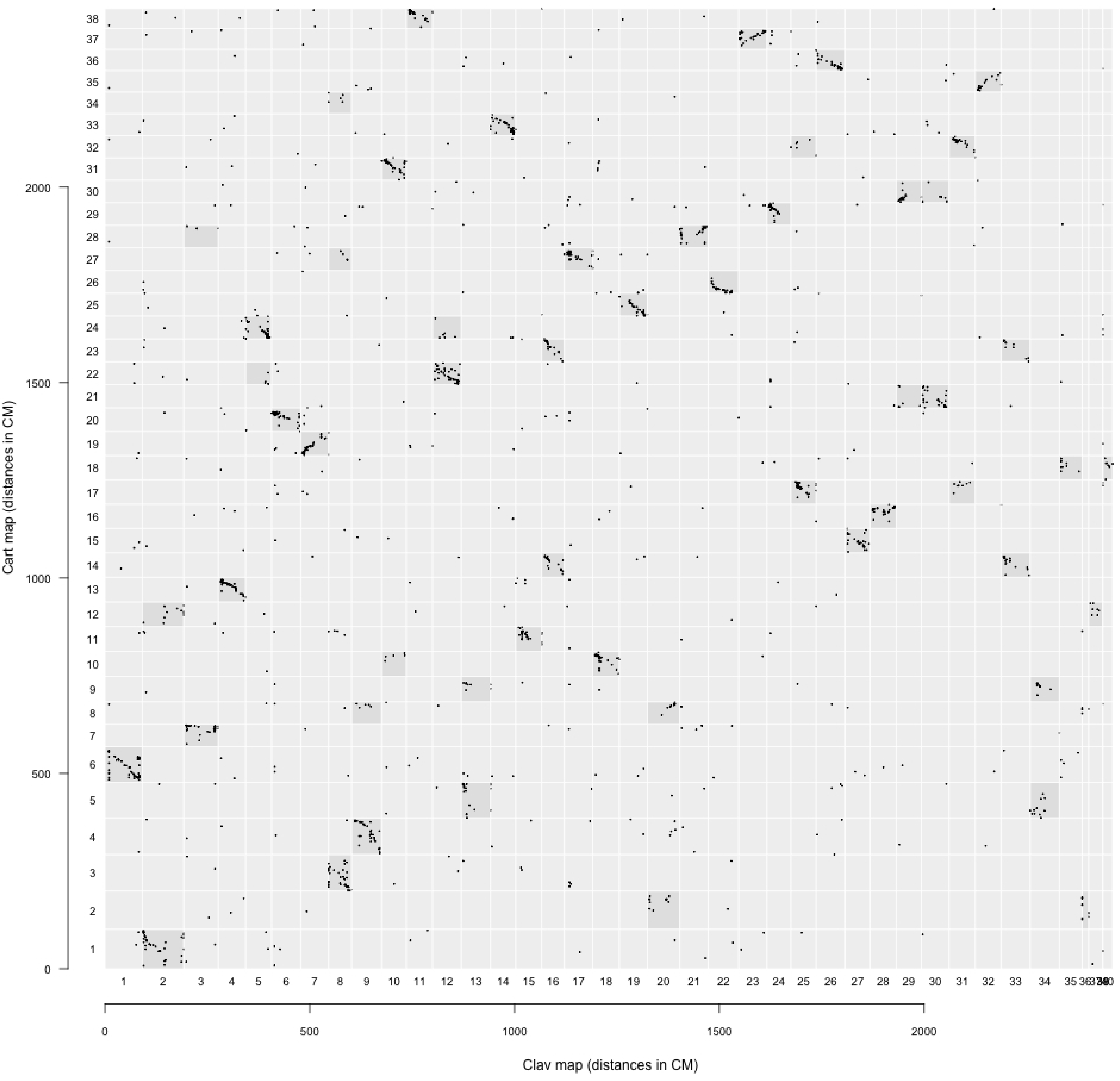

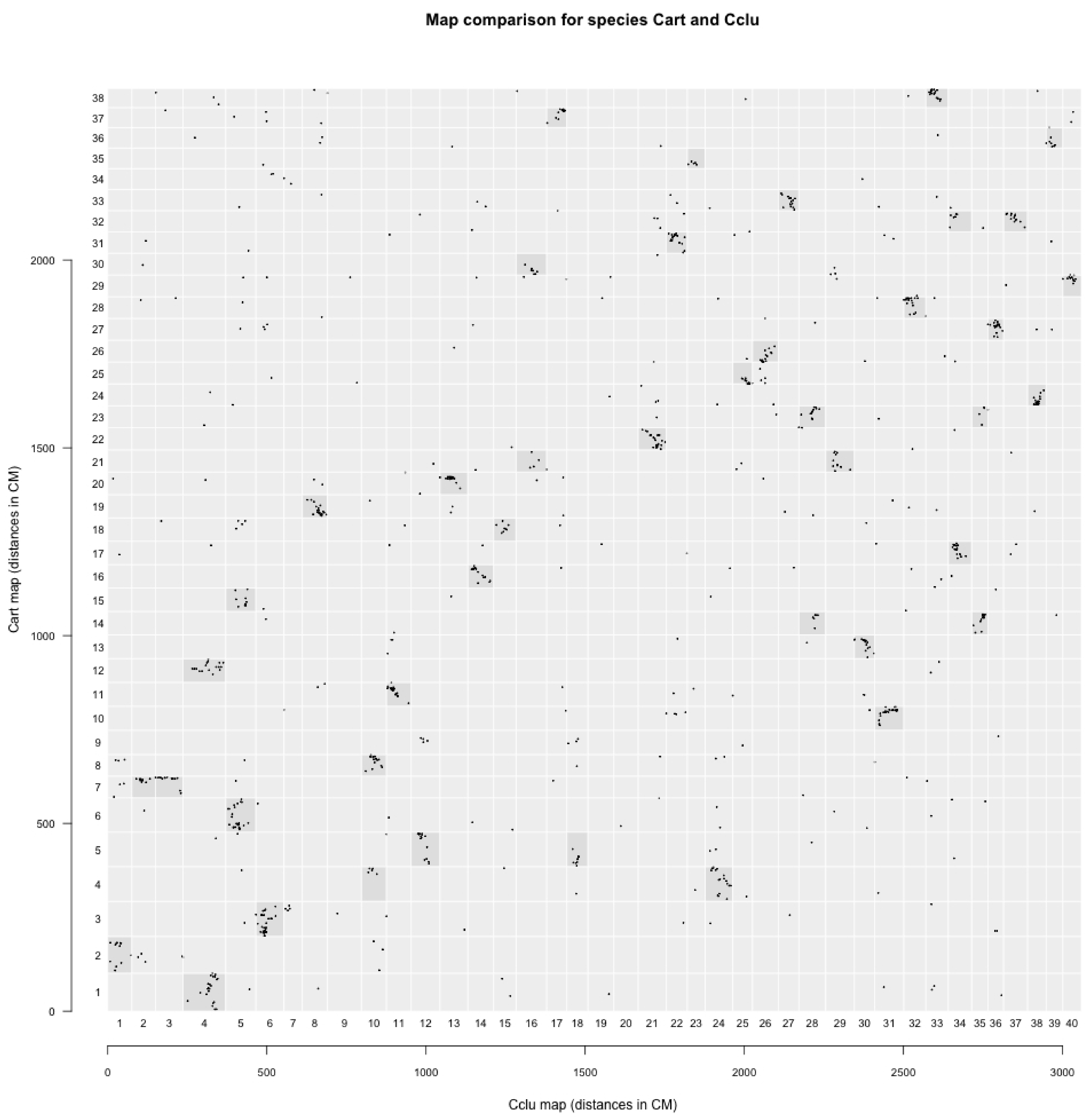

#### Map comparison for species Cart and Sfon

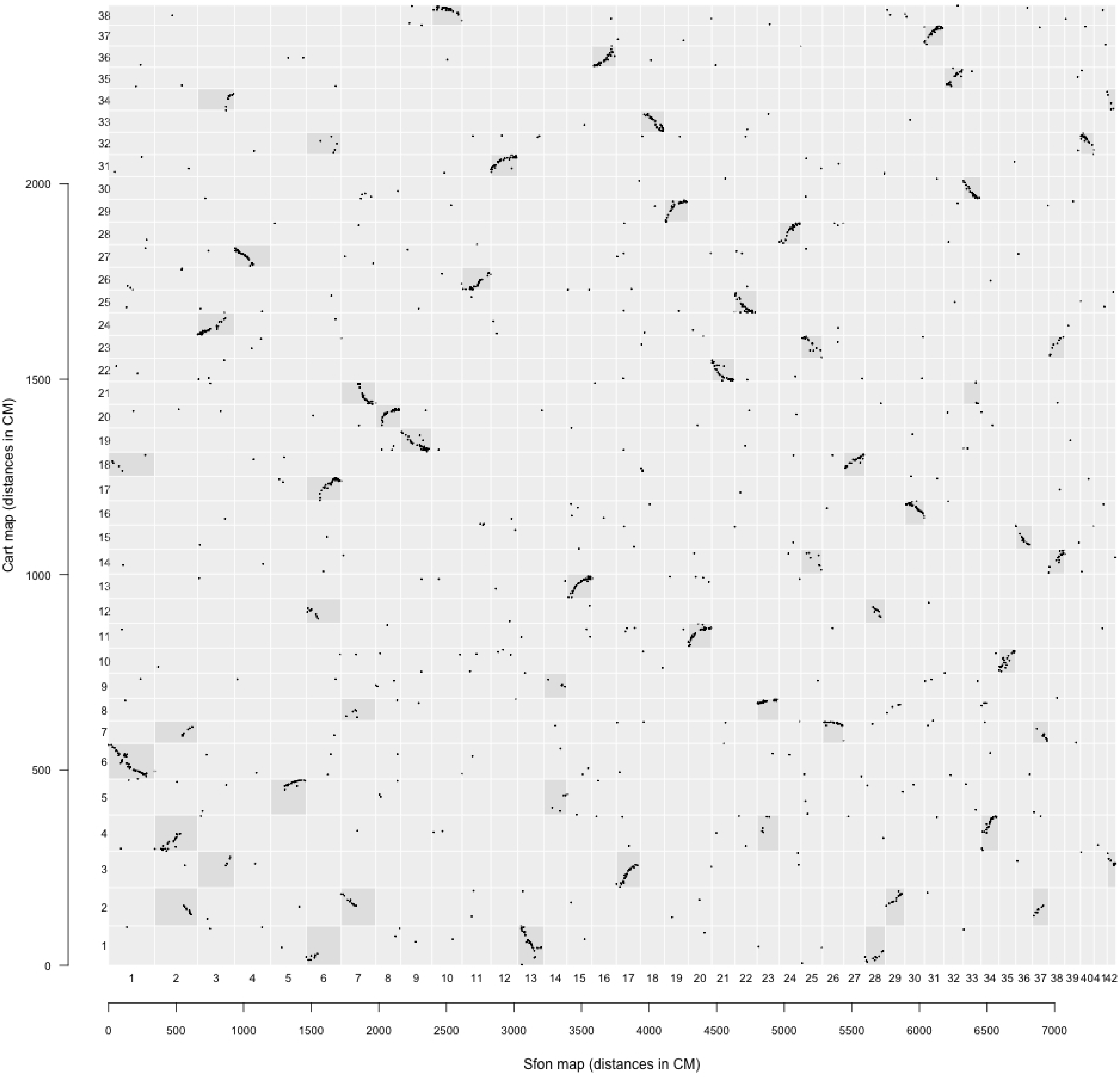

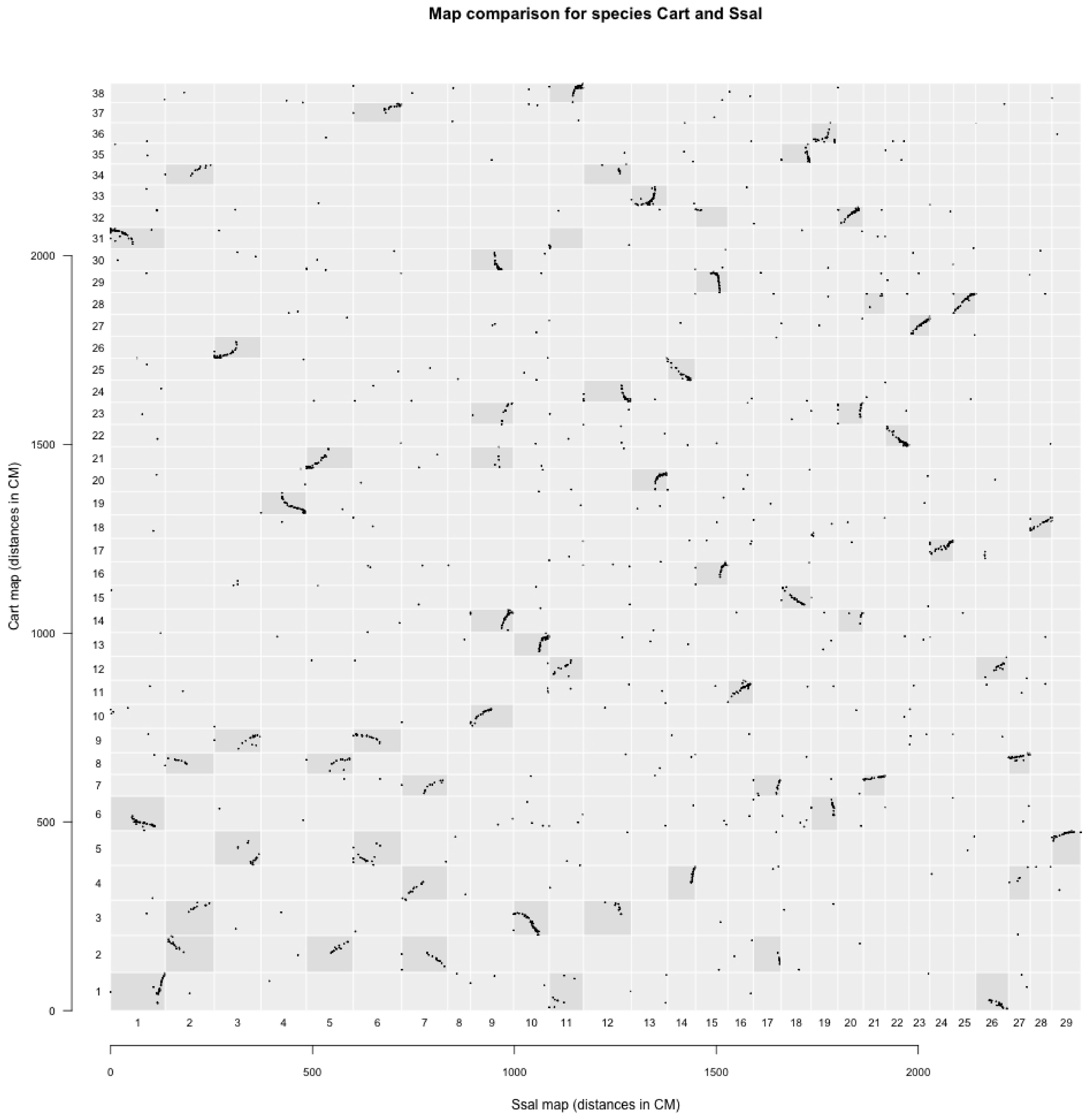

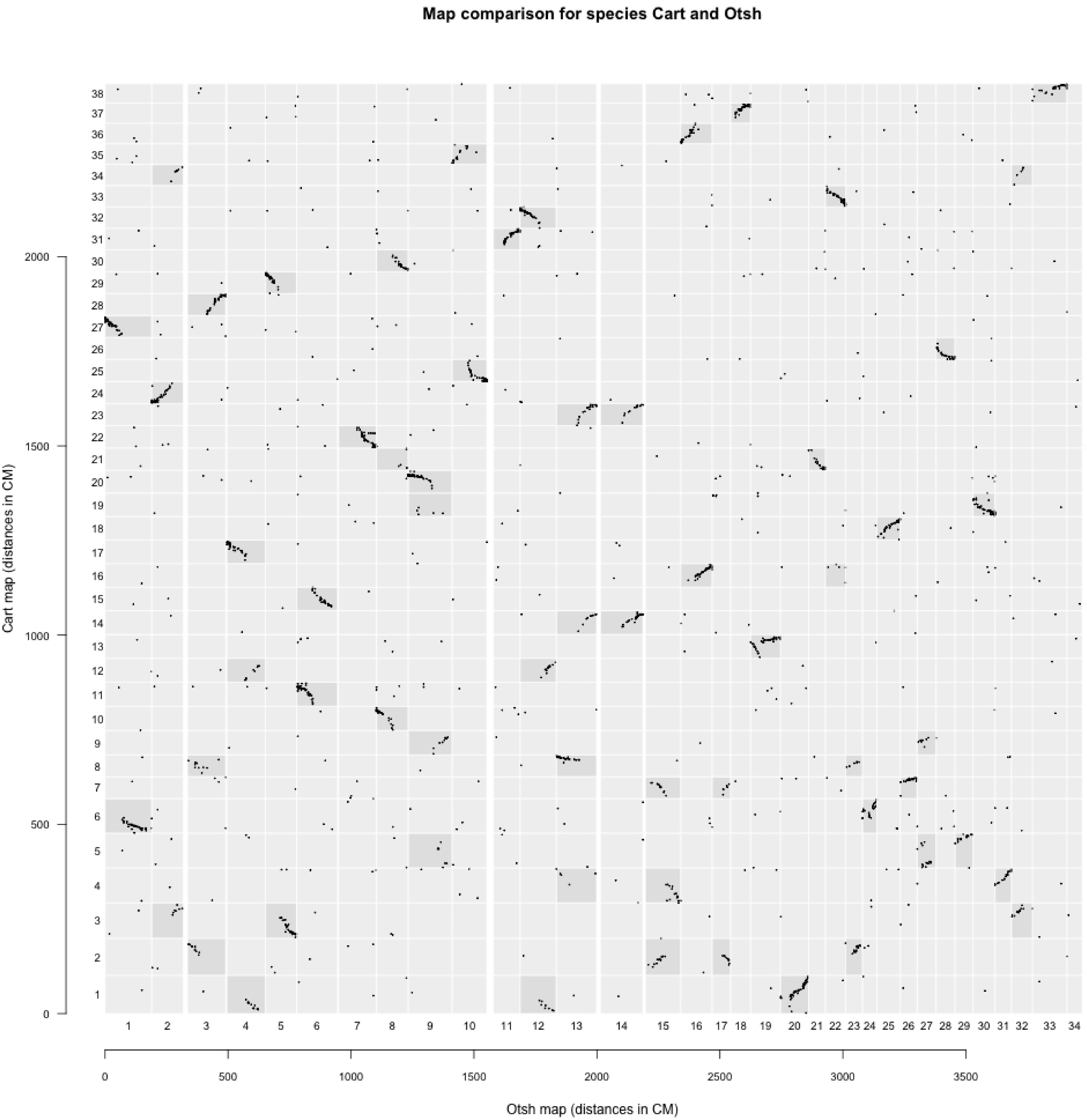

### Supplemental File 9

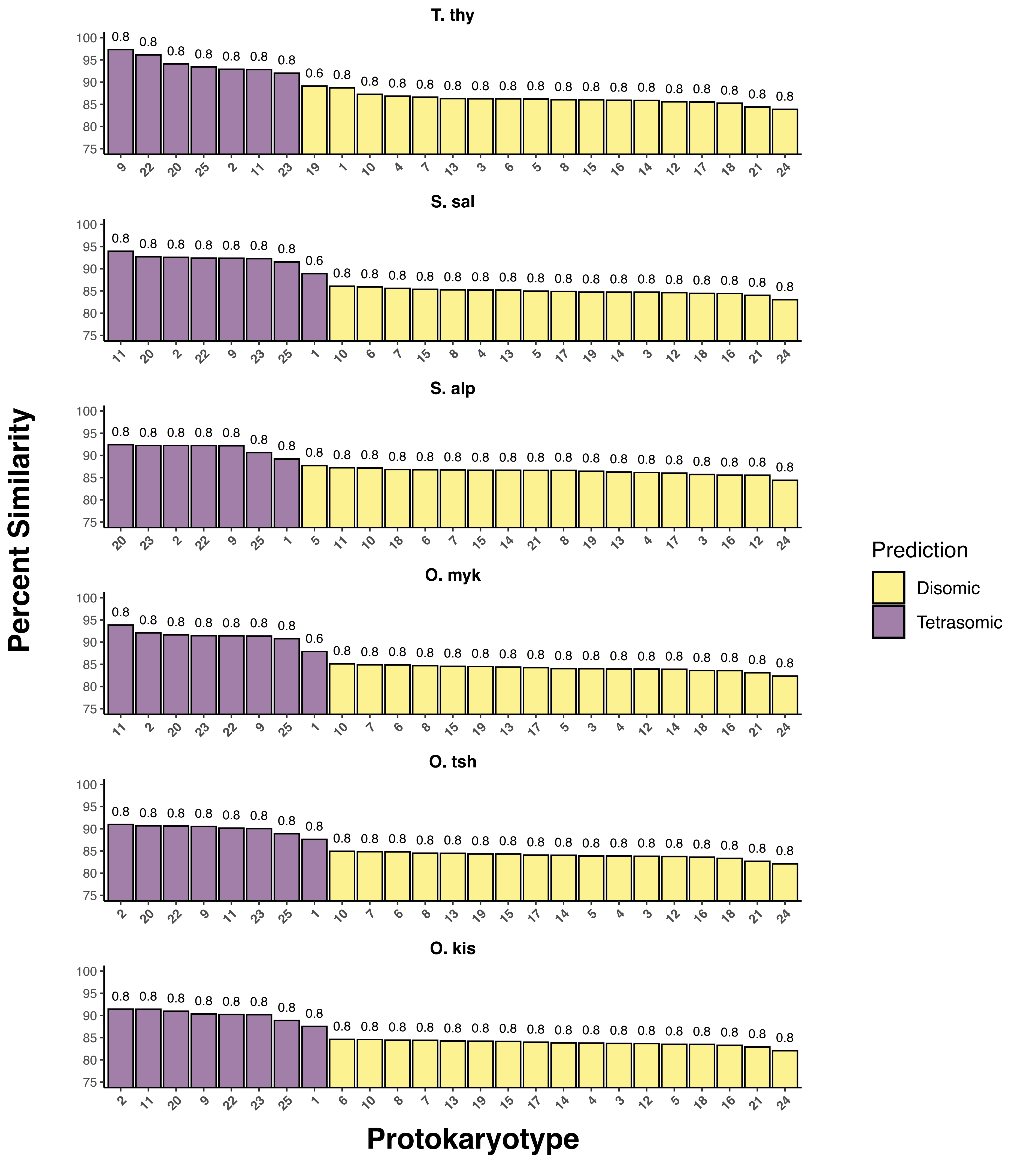
